# Cancer-Associated USP28 Missense Mutations Disrupt 53BP1 Interaction and p53 Stabilization

**DOI:** 10.1101/2025.05.20.655034

**Authors:** Hazrat Belal, Esther Feng Ying Ng, Midori Ohta, Franz Meitinger

## Abstract

Cellular stress response pathways are essential for genome stability and are frequently dysregulated in cancer. Following mitotic stress, the ubiquitin-specific protease 28 (USP28) and the p53-binding protein 1 (53BP1) form a stable, heritable complex to stabilize the tumor suppressor p53, triggering cell cycle arrest or apoptosis. Here we determine the mechanism by which USP28 stabilizes p53 and show that USP28 is required not only for an efficient stress response but also for maintaining basal p53 levels in some cancer cells. Loss of functional USP28 allows cells to evade mitotic stress and DNA damage responses in a manner that is specific to cell type and cancer context. We identify a prevalent, shorter USP28 isoform critical for p53 stabilization. Its C-terminal domain mediates PLK1-dependent binding to 53BP1, a dimerization-driven interaction necessary for mitotic stress memory, p53 stabilization, and cell cycle arrest. Cancer-associated missense mutations in this domain disrupt 53BP1 binding, impair nuclear localization, and destabilize USP28, compromising p53 stabilization. Notably, mutations in the 53BP1-binding domain occur more frequently in tumors than those in the catalytic domain, suggesting a potential role in cancer progression and implications for therapeutic strategies.

## Introduction

Mechanisms that regulate cell proliferation and genome stability are critical for tissue homeostasis. In response to cellular stress, diverse pathways converge on the tumor suppressor p53, leading to its stabilization and activation. This triggers cell cycle arrest or apoptosis—key barriers against genome instability and oncogenic transformation. The ubiquitin-specific protease 28 (USP28) has emerged as a key player in this process. While USP28 is known to promote tumor growth by stabilizing MYC in certain cancers ^1–5^, it has also been implicated in tumor suppression through p53 stabilization under stress conditions ^6–10^. These seemingly paradoxical roles raise important questions about the context-specific functions of USP28 in cancer. Notably, studies linking USP28 to MYC stabilization have primarily focused on p53-deficient cancer cells, suggesting that USP28’s role may shift depending on p53 status. Here, we define the molecular mechanism that engages USP28 in stress responses across normal and p53-wildtype cancer cells.

Systematic analysis of the cancer-associated mutations have identified USP28 and the p53-binding protein 1 (53BP1) as tumor suppressors, though the underlying mechanisms remained unclear ^11^. In 2016, three independent studies revealed that both proteins detect mitotic stress-induced prolonged mitosis, stabilizing p53 to trigger cell cycle arrest or apoptosis ^7–9^. This pathway, termed the “mitotic stopwatch” or “mitotic surveillance” pathway, protects cells from mitotic stress that could lead to chromosome missegregation and genome instability—hallmarks of cancer ^6–9,12–14^. While both USP28 and 53BP1 have been observed at DNA damage sites, USP28 does not directly participate in DNA repair but may contribute to local p53 activation ^10,15–17^. Notably, USP28’s roles in mitotic stress response and DNA damage response appear to be independent ^7-9,13,15,18,19^.

Recent work has shed light on the molecular basis of the mitotic stopwatch pathway. During prolonged mitosis, the mitotic kinase PLK1 facilitates the assembly of a USP28–53BP1– p53 complex ^13^. 53BP1 acts as a scaffold, recruiting USP28 and p53 via distinct domains, while PLK1-mediated phosphorylation of 53BP1 promotes p53 binding. However, the mechanisms regulating USP28’s engagement in this process remain unclear. After mitotic stress, this complex persists in daughter cells, stabilizing p53 and activating its transcriptional function. In the G1 phase, p53 induces the expression of p21^CDKN1A^, inhibiting CDK4/6 to prevent cell cycle progression. Alternatively, p53 can trigger apoptosis, as observed in human embryonic stem cells and mouse embryo development ^13,20^.

Growing evidence highlights the mitotic stopwatch’s role in maintaining tissue integrity during development ^19–26^. Centrosomal defects, which prolong mitosis in the developing mouse embryo, have been shown to induce p53-mediated apoptosis ^20^. Mutations in centrosomal genes frequently cause primary microcephaly, a condition associated with excessive mitotic stress in neuronal progenitors ^19,27^. Deletion of USP28 or 53BP1 alleviates these developmental defects, underscoring their critical role in responding to mitotic stress. Similar effects have been observed in epidermal stratification and embryonic lung and kidney development, further supporting the importance of USP28 in safeguarding tissue integrity ^20,22,24–26^. Considering the importance of USP28, it is crucial to understand the molecular modules required for the implementation of mitotic stress response.

Structurally, USP28 is a unique ubiquitin-specific protease (USP) containing an internal dimerization arm ^28,29^. Its USP domain is essential for both its tumor-suppressive and oncogenic functions ^5,7^. Two N-terminal ubiquitin-binding domains (UIM, UBA) have been proposed to mediate substrate interactions, though their role in substrate specificity remains unclear ^30^. USP28 dimerization is critical for its function ^28,29^, but recent findings suggest that DNA damage induces ATM-dependent phosphorylation, locking USP28 in a monomeric state that promotes MYC stabilization and genome instability potentially contributing to tumorigenesis by counteracting SCF^FBW^^7^-mediated MYC degradation ^3^. The function of the C-terminal region, which comprises nearly 40% of the protein, remains unknown. How USP28 coordinates its tumor-suppressive and oncogenic roles is a fundamental open question

In this study, we elucidate the molecular mechanism by which USP28 regulates p53 stability. We demonstrate that cancer-associated mutations in the C-terminal region of USP28 disrupt its dimerization- and PLK1-dependent interaction with 53BP1, selectively impairing its ability to stabilize p53 and coordinate stress responses. Notably, we find that a subset of cancer cells relies on USP28 to maintain basal p53 levels. As a result, USP28-deficient cancer cells exhibit attenuated responses not only to mitotic stress but also to DNA damage. Thus, USP28 missense mutations enable continued proliferation under stress conditions, potentially promoting genomic instability and driving tumor progression.

## Results

### USP28 is crucial for p53 stabilization in a cell type-specific manner

USP28 has been shown to stabilize the oncogene MYC to drive cell proliferation in response to DNA damage and the tumor suppressor p53 to cease cell proliferation in response to mitotic stress-induced prolonged mitosis or DNA damage ^6–9,13,17,18^. Considering these reported opposing functions of USP28, we asked the question if USP28 knockout confers an advantage or disadvantage to cancer cells that experience mitotic stress or DNA damage, specifically in untransformed and cancer-derived p53-wildtype cell lines. To do so, we developed a cell proliferation competition assay (**Fig. 1A**). A mixture of wildtype and *USP28Δ* cells were treated with the anti-mitotic inhibitor for PLK4 (PLK4i, centrinone, 150 nM) or the DNA damaging compound Doxorubicin (DXR, 10 nM). PLK4i induces mitotic stress by prolonging mitosis for 60-150 min and Doxorubicin introduces DNA strand breaks ^9,31^. The concentrations of both drugs were titrated to obtain similar effects on cell proliferation in p53-wildtype and p53-depleted hTERT RPE-1 (RPE1) cells (**Fig. 1B**). After eight days of treatment, the percentage of knockout cells was compared to wildtype cells. An enrichment of *USP28Δ* cells suggests that p53-dependent cell cycle arrest is the dominant mechanism, whereas a decrease indicates that MYC-dependent promotion of cell cycle progression prevails. We also performed this assay for 53BP1, which is required for USP28-dependent p53 stabilization. For this experiment, we selected one non-transformed and eleven cancer cell lines that express wildtype p53 and exhibit comparable proliferation rates. We found that the deletion of *USP28* or *TP53BP1* significantly reduced the sensitivity of several cancer cells to PLK4i treatment, mirroring the response observed in untransformed RPE1 cells (**Fig. 1C**). In contrast, *USP28Δ* had a less pronounced effect, and *TP53BP1Δ* had no effect on the sensitivity to DXR-induced DNA damage. The later observation could be explained by the independent role of 53BP1 in DNA repair ^32^. To further investigate the role of USP28, we analyzed the impact of mitotic stress and DNA damage on the stability of p53 and MYC. Both stress conditions led to p53 stabilization, p21 expression and MYC downregulation in a USP28-dependent manner (**Fig. 1D**). However, while MYC downregulation still occurred in *USP28Δ* cells following DNA damage, it was less pronounced than in wildtype cells. A similar trend was observed for RPE1 cells that harbor a mutation in 53BP1 (G1560K) that impairs binding to USP28, suggesting that the observed effects rely on this interaction. Surprisingly, we found that *USP28* deletion reduced the p53 amount in unstressed cancer cells (A549 and U2OS) (**Fig. 1D, S1A)**. Probably due to the reduced p53 levels, *USP28Δ* A549 cells failed to respond to both mitotic stress and DNA damage (**Fig. 1D**). In contrast to previous studies^3–5^, *USP28* deletion had no effect on the half-life of MYC in the four tested p53 wildtype cell lines **(Fig. S1B-F)**.

**Figure 1:**
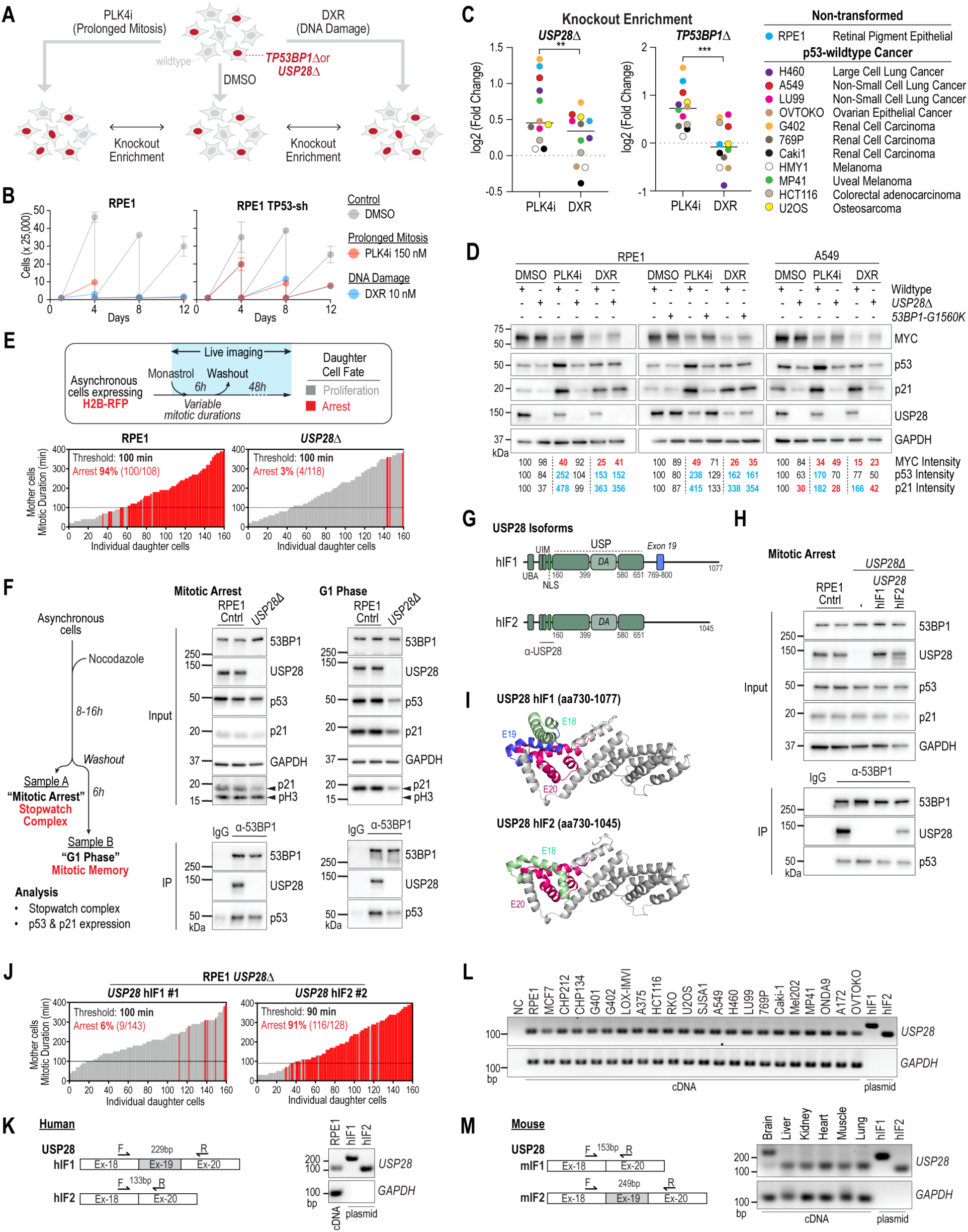
Isoform-specific requirement of USP28 for mitotic stress-induced stabilization of p53 **(A)** Schematic of the competition assay used to assess USP28- and 53BP1-dependent sensitivity to drugs. Wildtype and gene deletion mutants were mixed and treated with PLK4 inhibitor (PLK4i, 150 nM) or Doxorubicin (DXR, 10 nM). After 8 days, knockout cell abundance was measured by sequencing. Enrichment of knockout cells indicates gene-dependent drug sensitivity. **(B)** Proliferation assay of RPE1 and RPE1 TP53-sh cells treated with PLK4i or DXR. **(C)** Enrichment of *USP28Δ* and *TP53BP1Δ* cells following 8-day PLK4i or DXR treatment. **(D)** Immunoblotting of RPE1 and A549 cells with indicated genotypes after 4-day drug exposure. Blue and red numbers indicate samples with p53/p21 enrichment (>1.5× DMSO) and MYC downregulation (<0.5× DMSO), respectively. GAPDH, loading control. **(E)** Imaging-based assay evaluating daughter cell fate after prolonged mitosis in wildtype and *USP28Δ* RPE1 cells. Cells underwent transient mitotic arrest via Monastrol, followed by washout and imaging for 48 h. Each bar represents a daughter cell; grey indicates division, red indicates arrest. Bar height corresponds to mother cell mitotic duration. The arrest threshold is defined as the mitotic duration where >50% of daughters arrest. **(F)** 53BP1 immunoprecipitation from wildtype and *USP28Δ* RPE1 cells arrested in mitosis with Nocodazole (8–16 h) or released into G1. “Mitotic Arrest” samples were used to detect stopwatch complexes (53BP1–USP28–p53); “G1 Phase” samples assessed p53 activation. IP, immunoprecipitates. Inputs, soluble supernatants. GAPDH, loading control. pH3, mitosis marker. **(G)** Schematic of human USP28 isoforms. Exon 19 is present in USP28^hIF1^ but not USP28^hIF2^. hIF1, human isoform 1; hIF2, human isoform 2; UBA, Ubiquitin-associated domain; UIM, Ubiquitin-interacting motif; NLS, Nuclear localization sequence; USP, Ubiquitin-specific protease; DA, Dimerization arm. The epitope of the antibody used for USP28 recognition (α-USP28) is shown. **(H)** 53BP1 immunoprecipitation from mitotically arrested *USP28Δ* cells expressing USP28^hIF1^ or USP28^hIF2^, assessing 53BP1 binding. **(I)** AlphaFold model of USP28 C-terminus. Isoform-specific structural differences include loss or shift of helices in USP28^hIF2^. **(J)** Imaging assay as in (E), showing USP28^hIF2^-expressing cells arrest after prolonged mitosis, unlike hIF1-expressing cells. **(K–M)** RT-PCR analysis of exon 19 expression. Exon 19 is not detected in RPE1 cells (K), across 21 cancer lines (L), or mouse tissues (M). Controls: cDNAs for hIF1 and hIF2. GAPDH, loading control.

Collectively, these findings suggest that USP28, through its interaction with 53BP1, plays a crucial role in mediating the cellular response to mitotic stress. In certain cancer cells, USP28 is essential for maintaining baseline p53 levels under unstressed conditions. Consequently, its deletion disrupts both the mitotic stress response and DNA damage response.

### USP28 Isoform-specific response to mitotic stress

USP28 is frequently mutated in cancer ^11^. To recapitulate cancer-associated mutations and determine their consequences on USP28-mediated control of p53, we established a system in RPE1 cells where we can replace wildtype USP28 with mutant transgenes. We first assessed the USP28-dependent sensitivity of RPE1 cells to mitotic stress (extended mitosis) using a live-cell imaging approach (**Fig. 1E**) ^9,13^. Cells were labeled with H2B-RFP and transiently treated with the KIF11^Eg^^5^ inhibitor Monastrol, which induces a reversible mitotic arrest ^18^. During treatment, cells were monitored and tracked for six hours using fluorescence imaging. Each cell in the asynchronous population entered mitosis at different time points and remained arrested in mitosis. Following Monastrol washout, cells exited mitosis and completed cell division. This approach gave rise to daughter cells whose mother cells had different mitotic lengths. Daughter cells were then tracked for an additional 48 hours to assess their fate.

As previously reported, the progeny of cells that spent over ninety minutes in mitosis exhibited a stable cell cycle arrest (**Fig. 1E**) ^9,13,18^. However, the deletion of *USP28* rendered cells less sensitive to mitotic stress, allowing them to continue proliferating despite a parental mitotic duration exceeding ninety minutes (**Fig. 1E**). This sensitivity to mitotic stress is dependent on the formation of a complex involving 53BP1, USP28, and p53 (**Fig. 1F, S2A)** ^13^. We observed that this complex remains stable following mitotic exit, facilitating p53 stabilization and subsequent p21 expression (**Fig. 1F**). In USP28-deleted cells, the response to prolonged mitosis was impaired, resulting in a failure to stabilize p53 and p21.

The canonical isoform *USP28*^hIF^^1^ (human isoform 1, NP_065937.1, NM_020886.4) encodes a 1077-amino acid protein (**Fig. 1G; S2B)**. To investigate its role, we generated a *USP28* knockout cell line ectopically expressing *USP28*^hIF^^1^ but found that the longer isoform USP28^hIF1^ did not interact with 53BP1 in mitotically arrested cells (**Fig. 1H**). In contrast, the shorter isoform, USP28^hIF2^ (human isoform 2, NP_001333187.1, NM_001346258.2), successfully interacted with 53BP1, suggesting that isoform 2, rather than isoform 1, mediates the mitotic stress response. Notably, isoform 2 lacks exon 19, which encodes a 32-amino acid sequence in the C-terminal domain (**Fig. 1G; S2B)**. Structural predictions from AlphaFold with high-confidence modeling indicate that exon 19 forms an additional alpha helix in isoform 1, which may obstruct the interaction surface for 53BP1 (**Fig. 1I; S2C)**. We established single clones that express similar amount of USP28^hIF1^ and USP28^hIF2^ compared to parental RPE1 cells **(Fig. S2D)**. In line with the interaction capability, only USP28^hIF2^ but not USP28^hIF1^ expressing cells induced cell arrest following prolonged mitosis (**Fig. 1J**).

Prompted by these findings, we analyzed the expression levels of both isoforms in RPE1 cells and found that USP28^hIF2^ is predominantly expressed (**Fig. 1K**). To explore whether isoform-specific expression is consistent across different cell types, we extended our analysis to 22 p53-wildtype cancer cell lines from 10 tissue origins **(Fig. S2E)**. In all cell lines examined, the short isoform USP28^hIF2^ was the predominant transcript (**Fig. 1L**). We also analyzed the expression of USP28 isoforms in six mouse tissues, where the shorter isoform is designated as isoform 1 (USP28^mIF1^, NP_780691.2, NM_175482.3), which exhibits similarity to human isoform 2 and the longer isoform is designated as isoform 2 (USP28^mIF2^, NP_001346668.1, NP_780691.2), resembling human isoform 1 **(Fig. S2B)**. In most mouse tissues, the shorter isoform was also dominant, except in the brain, where a higher proportion of isoforms containing exon 19 was detected (**Fig. 1M**). Sequencing confirmed that this band corresponded to exon 19 of the longer isoform of USP28 **(Fig. S2F)**. We subsequently tested additional human and mouse cell lines originating from neuronal tissues; however, none expressed notable levels of the longer isoform **(Fig. S2G, H)**.

In conclusion, our findings demonstrate that the short isoform USP28^hIF2^ is essential for the mitotic stress response, while the long isoform USP28^hIF1^ is unable to fulfill this role. This suggests an isoform-specific regulatory mechanism for USP28 in the mitotic stress response and highlights the importance of the USP28 C-terminus on tumor-suppressor function.

### Identification of mutations in USP28 that desensitize cells to mitotic stress

The USP domain of USP28 is required for p53 and MYC stabilization ^5,7^. The motifs and domains of USP28 that are specifically required for p53 stabilization are not known. Thus, we set out to determine the functions of the regions that are N-terminal and C-terminal of the USP domain of USP28. To do so we took advantage from a surprising observation. When we expressed USP28^hIF2^ in *USP28Δ* RPE1 cells, we noted that 16 out of the 34 isolated cell clones carried mutations in USP28^hIF2^ (**Fig. 2A-B; S3A)**. Each mutation was unique to a specific clone, ruling out the possibility of mutations originating from the lentiviral construct used to express the transgene. Five clones contained frameshift mutations, and two carried nonsense mutations with premature stop codons. Since USP28 overexpression is toxic, we reasoned that all the observed mutations specifically impair p53 stabilization (**Fig. 2B; S3A)** ^7^. The identified missense mutations were located in the ubiquitin-specific protease domain (USP; aa149-399 and aa480-650; 5 clones), the dimerization arm (DA; aa400-579; 3 clones), and the C-terminus (aa651-1045; 5 clones). We selected six clones with one or two mutations in USP28^hIF2^ and assessed their expression levels. For comparison, we also generated a clone expressing a catalytically inactive mutant of USP28^hIF2^ (C171A) ^7^. All tested USP28^hIF2^ mutants were expressed at levels similar to or higher than endogenous USP28 **(Fig. S3B)**.

**Figure 2:**
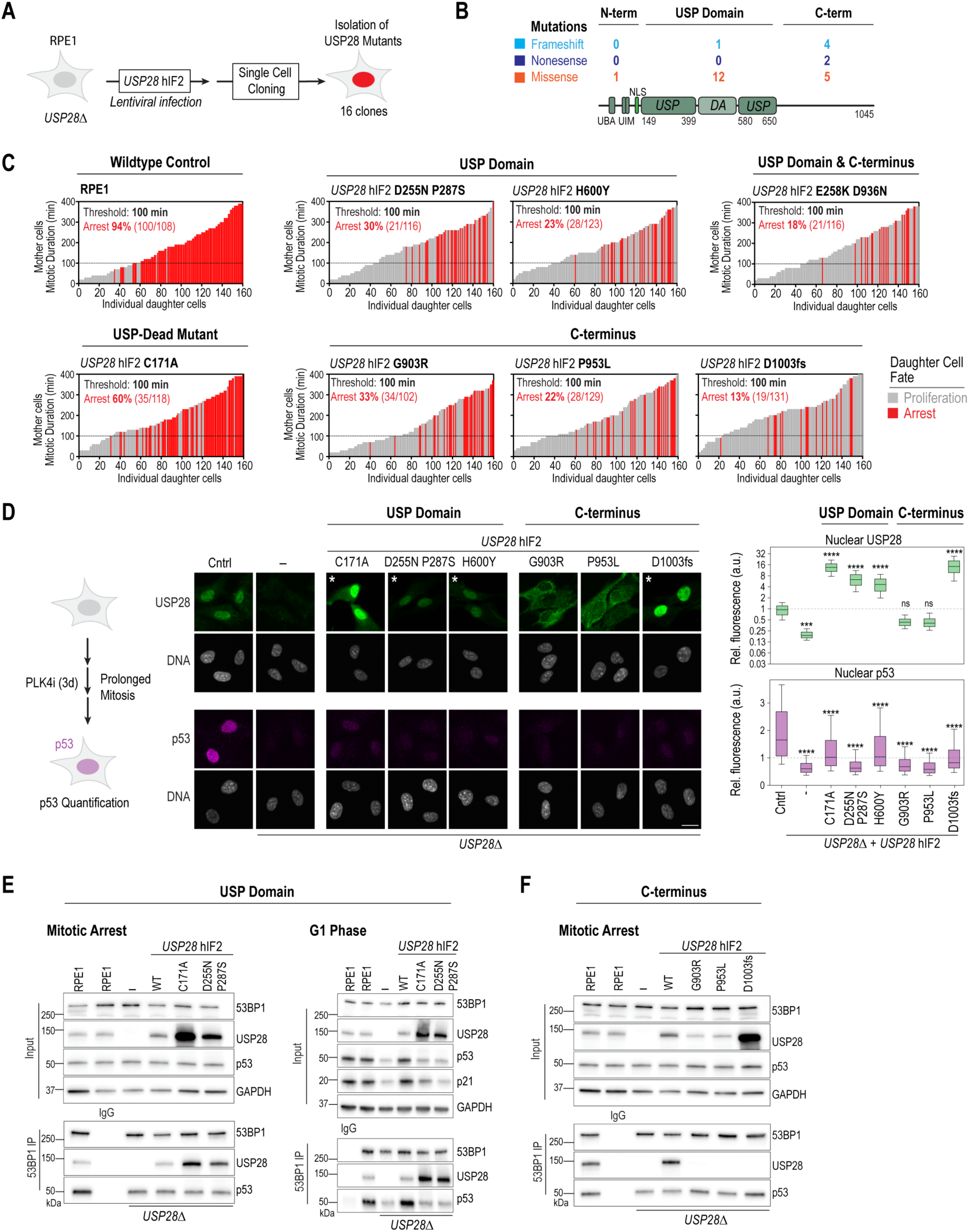
Identification of mutations in USP28 that desensitize cells to mitotic stress (A) Schematic showing isolation of cell clones expressing mutant variants of the *USP28^hIF2^* transgene. Mutations occurred spontaneously without applied stress. (B) Schematic of USP28 mutations identified in clones from (A), including missense, nonsense, and frameshift changes. Mutations were categorized by location: N-term (aa 1–149), USP domain (aa 149–650), and C-term (aa 651–1045). See also Figure S3A. (C) Imaging assay (as in Figure 1E) showing that USP28-mutant clones display impaired response to prolonged mitosis. The RPE1 control graph is reused from Figure 1E. A USP-dead control (C171A) is included. Graphs are grouped by mutation region as in (B). (D) Microscopy-based analysis of USP28 and p53 expression in RPE1, *USP28Δ*, and USP28Δ cells expressing *USP28*^hIF2^ mutants. USP28 panels marked with an asterisk were displayed at 5× lower intensity (see Figure S3C for unadjusted images). Nuclei were stained with Hoechst 33342. Quantification of nuclear USP28 and p53 levels is shown (upper and lower right graphs, respectively). Cells were treated with PLK4i for 3 days to induce prolonged mitosis. Data are box-and-whisker plots normalized to untreated RPE1 cells. Mean and 10–90 percentiles shown. Statistical analysis by one-way ANOVA (*P < 0.05; **P < 0.01; ***P < 0.001; ****P < 0.0001). n = 1,000 cells/condition. Scale bar: 10 µm. **(E–F)** Co-immunoprecipitation analysis of 53BP1 complexes in mitotically arrested and G1 phase cells, assessing USP28 and p53 binding and stabilization. Experimental setup as in Figure 1F; *USP28*^hIF2^ wildtype and mutant clones were isolated as shown in (A). (E) Analysis of wildtype *USP28*^hIF2^ and two USP domain mutants (C171A, D255N/P287S). (F) Analysis of wildtype *USP28*^hIF2^ and three C-terminal mutants (G903R, P953L, D1003fs). Unmodified RPE1 cells served as controls. Inputs, soluble fractions. IP, immunoprecipitates. GAPDH, loading control.

To assess the functionality of the USP28^hIF2^ mutants, cells were labeled with H2B-RFP and analyzed for sensitivity to prolonged mitosis using live imaging and single-cell tracking (**Fig. 1E; 2C)**. In wildtype RPE1 cells, 94% of cells that experienced mitotic durations exceeding 90 minutes underwent arrest. In contrast, the C171A control mutant and the USP28^hIF2^ mutants displayed significantly reduced sensitivity to prolonged mitosis. Among the mutants, only 13% to 33% of cells were arrested following extended mitosis (> 90 minutes), indicating that these mutations impair USP28’s role in the mitotic stress response.

While clones with mutations in the ubiquitin-specific protease domain (C171A; D255N P287S; H600Y) and one clone with a frameshift mutation in the C-terminus (clone 16, D1003fs) exhibited nuclear localization similar to wildtype USP28^hIF2^, two clones with mutations in the C-terminus (G903A and P953L) failed to localize to the nucleus (**Fig. 2D; S3C)**. USP28^hIF2^ has a predicted nuclear localization sequence (NLS) in the N-terminus (aa135-145; **Fig. 2B; S3A**), but this sequence does not explain the altered cytoplasmic localization observed in the G903A and P953L mutants (**Fig. 2B, D; S3A, C)**. Notably, the expression levels of all tested mutants, except G903A and P953L, were 5- to 20-fold higher than in wildtype RPE1 cells (**Fig. 2D; S3B, C)**, suggesting that the mutants fail to stabilize p53 following an extended mitotic duration even though the expression level is significantly increased. To assess p53 activation, we treated cells with the PLK4 inhibitor to prolong mitosis and measured stabilized p53 in the nucleus three days after the start of the treatment (**Fig. 2D**). We found that all mutant clones failed to sufficiently stabilize p53 after prolonged mitosis (**Fig. 2D**).

To investigate the molecular defects in the USP28^hIF2^ mutants, we assessed their ability to form a complex with 53BP1 during prolonged mitosis (**Fig. 2E, F**). Mutations in the USP domain (C171A; D255N P287S) did not affect USP28^hIF2^ binding to 53BP1 (**Fig. 2E**). However, after release from mitotic arrest into the G1 phase, the complex remained stable but failed to stabilize p53 in the tested USP mutants (**Fig. 2E**). This suggests that the mutations impair USP28’s deubiquitinase activity, which is essential for p53 stabilization ^7^. Strikingly, the three mutants with distinct mutations in the C-terminal domain failed to interact with 53BP1, indicating that the C-terminus of USP28^hIF2^ is required for its interaction with 53BP1 and subsequent p53 stabilization after prolonged mitosis (**Fig. 2F**).

### The C-terminus and dimerization of USP28^hIF2^ are required for interaction with 53BP1

One of the isolated mutants that failed to interact with 53BP1 harbored a frameshift mutation near the C-terminus of the coding sequence (D1003fs). This mutation resulted in a 10-base pair deletion, leading to a C-terminally truncated protein (1007aa; full length is 1045aa) **(Fig. S3D)**. To identify the minimal region of USP28^hIF2^ required for interaction with 53BP1, we generated mutants expressing transgenes of shorter C-terminal truncations in *USP28Δ* RPE1 cells (**Fig. 3A**). We found that deletion of the last 13 amino acids was sufficient to disrupt the interaction with 53BP1, and this truncation failed to stabilize p53 following release from mitotic arrest (**Fig. 3A, B**). Taken together, our work identified four mutants that failed to interact with 53BP1 (**Fig. 3C**). The identified mutants have either a missense mutation (G903A, P953L) or a C-terminal truncation of 13 or more amino acids. The longer isoform USP28^hIF1^ has an additional alpha helix between amino acids 769 and 800, which impairs the interaction with 53BP1. These results indicate that the predicted C-terminal domain (aa651-1045) is required for the interaction with 53BP1.

**Figure 3:**
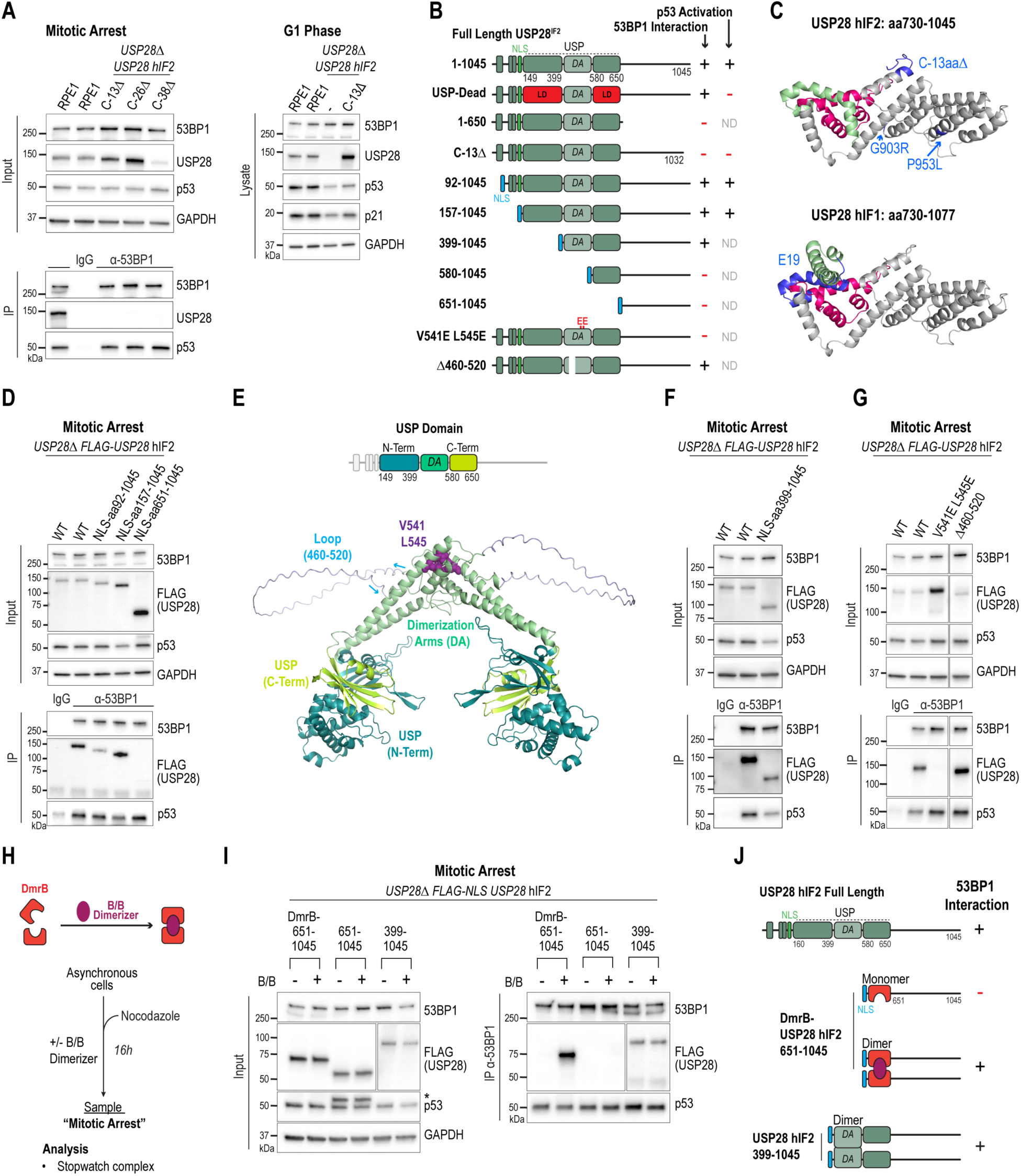
The C-terminus and dimerization of USP28^hIF2^ are required for interaction with 53BP1 **(A)** Analysis of 53BP1 immunoprecipitates and lysates from mitotically arrested cells to assess complex formation with USP28 and p53 (middle) and p53 stabilization in G1 phase (right). *USP28*^hIF2^ transgenes with C-terminal truncations (Δ13, Δ26, Δ38 aa) were expressed in *USP28Δ* RPE1 cells. Unmodified RPE1 served as control. Inputs, soluble supernatants. IP, immunoprecipitates. GAPDH, loading control. **(B)** Summary of USP28 mutants and their ability to bind 53BP1 and stabilize p53. Endogenous and nucleoplasmin-derived NLS sequences are indicated (green and blue). **(C)** AlphaFold-predicted structural model of the USP28 C-terminus, highlighting mutations in isoform 2 and the alpha helix of exon 19 in isoform 1 that interfere with 53BP1 binding. **(D)** Co-immunoprecipitation of 53BP1 with USP28 and p53 from mitotically arrested cells expressing N-terminal truncations of *USP28*^hIF2^ (aa92–1045, aa157–1045, aa651–1045). All constructs carry an N-terminal FLAG tag and nucleoplasmin NLS. GAPDH, loading control. **(E)** AlphaFold model of USP28 dimer highlighting the USP domain and dimerization arm (DA). Critical residues for dimerization (V541, L545) and the unstructured loop (aa460–520) are marked. Color codes match schematic above. **(F-G)** Immunoprecipitation analysis of 53BP1 complex formation in mitotically arrested cells expressing mutant *USP28*^hIF2^ transgenes. (F) A construct lacking the N-terminal region including part of the USP domain. (G) Constructs with dimerization-disrupting mutations (V541E, L545E) or loop deletion (Δ460–520). Wildtype *USP28*^hIF2^ serves as control. Blots in (G) are from the same membrane. IP, immunoprecipitated. GAPDH, loading control. **(H)** Schematic of an inducible dimerization domain. Asynchronous cells were treated for 16h with Nocodazole (100 ng/ml) and the B/B dimerizer (100 nM) to induce dimerization. Mitotic cells were harvested and analyzed by immunoprecipitation. **(I)** Co-immunoprecipitation to evaluate dimerization-dependent 53BP1–USP28–p53 complex formation. A C-terminal USP28 fragment (651–1045) fused to DmrB and FLAG was expressed in *USP28Δ* RPE1 cells. Controls included constructs lacking DmrB or containing the full dimerization arm (399–1045). See also Figure S4F. Asterisk in p53 blot indicates signal from earlier FLAG detection. IP, immunoprecipitated. GAPDH, loading control. **(J)** Summary of (H–I) results demonstrating that USP28 dimerization is essential for C-terminal interaction with 53BP1.

Next, we sought to determine the minimal region of USP28^hIF2^ required for 53BP1 binding. As expected, the N-terminal region (aa1-650), including the USP domain and dimerization arm, was unable to mediate 53BP1 interaction (**Fig. 3B; S4A)**. Surprisingly, the C-terminal region alone (aa651-1045) was also insufficient (**Fig. 3B; S4A-D)**. Forcing nuclear localization of the C-terminal fragment with a nuclear localization signal (NLS, from nucleoplasmin) did not enhance 53BP1 binding (**Fig. 3B, D; S4B-D)**. However, a larger fragment containing both the C-terminus and the ubiquitin-specific protease (USP) domain (aa157-1045) could interact with 53BP1 (**Fig. 3B, D**).

USP28 contains a unique USP domain interrupted by a dimerization arm (DA; aa400-579) ^28,29^. An AlphaFold model of the USP domain reveals both structured and unstructured regions within this domain (**Fig. 3E, S4E)**, closely resembling configurations of previously reported structures ^28,29^. Interestingly, a construct containing only the dimerization arm but lacking the N-terminal portion of the USP domain, was sufficient to interact with 53BP1 (**Fig. 3B, F**). To determine whether dimerization is necessary for this interaction, we tested a USP28 dimerization mutant, V541E L545E (**Fig. 3B, E, G**) ^28,29^. We found that this dimerization mutant was unable to interact with 53BP1. In contrast, an unstructured region (Δ460-520) within the dimerization arm was not necessary (**Fig. 3B, E, G**).

Our data suggest that 53BP1 either exclusively binds the C-terminus of dimerized USP28 or that the dimerization arms create an additional binding surface for 53BP1. To differentiate between these possibilities, we fused an inducible dimerization domain (DmrB) to the N-terminus of a C-terminal USP28 fragment (651-1045), which by itself was unable to interact with 53BP1. Strikingly, DmrB-induced dimerization of the C-terminal USP28 fragment drastically enhanced its interaction with 53BP1 (**Fig. 3H-J, S4F)**. A similar effect was observed with a slightly longer C-terminal fragment (580-1045) **(Fig. S4G-I)**. These results demonstrate that 53BP1 interacts exclusively with the dimerized form of the USP28^hIF2^ C-terminus.

### PLK1-mediated USP28 C-terminus interaction with 53BP1 underlies mitotic memory

The mitotic stress response relies on the transmission of mitotic stopwatch complexes, formed during prolonged mitosis, to progeny, a process known as mitotic memory ^13^. This intrinsic mechanism is driven by PLK1-mediated formation of the mitotic stopwatch complex (**Fig. 4A**). We hypothesized that the C-terminus of USP28 plays a critical role in mediating mitotic memory. Supporting this hypothesis, we found that the interaction between 53BP1 and the synthetically dimerized C-terminus of USP28 is PLK1-dependent (**Fig. 4B**). Although PLK1 is active only during mitosis, this complex remains stable after cells are released from prolonged mitotic arrest into the G1-phase (**Fig. 4C**). Thus, the intrinsic function of the C-terminus of USP28 is to mediate mitotic memory through its interaction with 53BP1.

**Figure 4:**
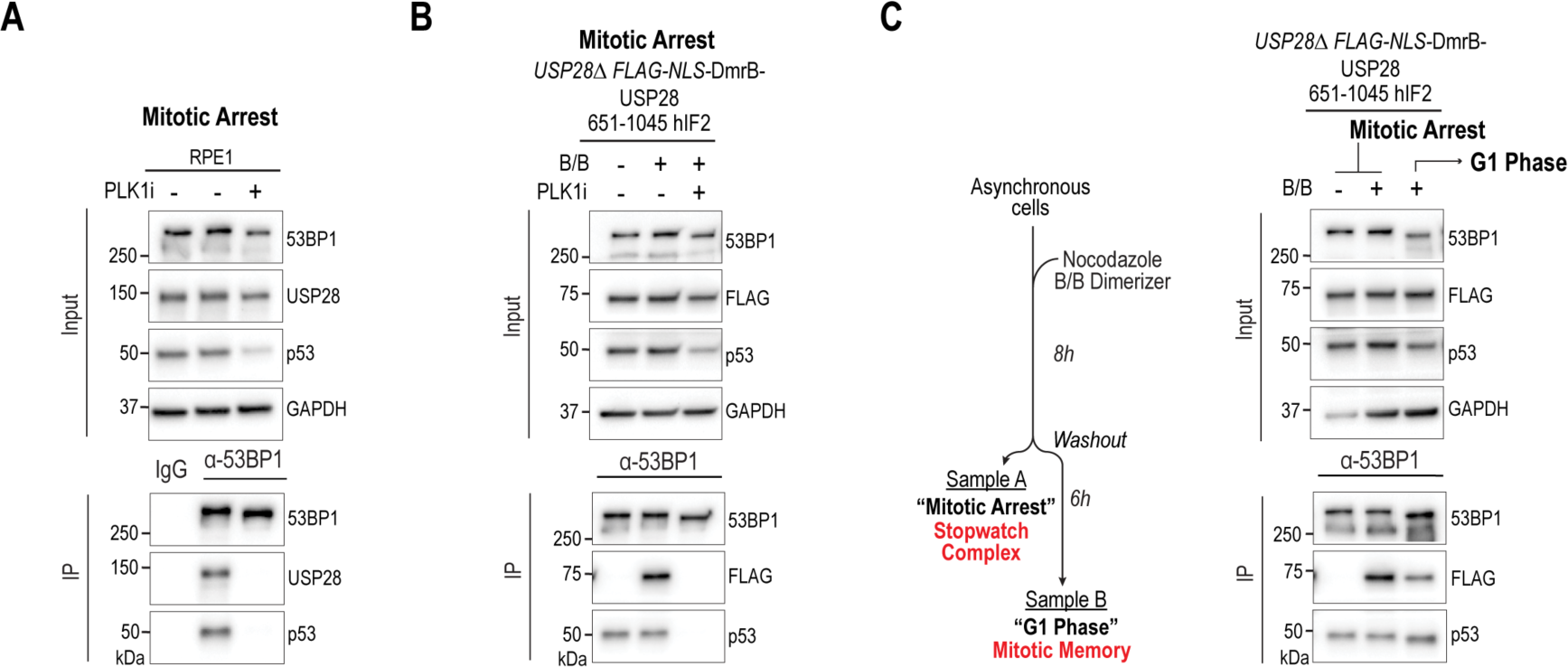
The stable interaction between C-terminus of USP28^hIF2^ and 53BP1 mediates mitotic memory **(A)** Immunoprecipitation analysis of 53BP1 in cells treated with or without PLK1 inhibitor to assess the role of PLK1 kinase activity in its interaction with USP28. **(B)** Immunoprecipitation analysis of 53BP1 in cells expressing a synthetic dimerized USP28 C-terminal construct, performed with or without PLK1 inhibition. **(C)** Immunoprecipitation analysis of 53BP1 after release from mitotic arrest into G1 phase, used to evaluate the stability of its interaction with the synthetic dimerized USP28 C-terminus. IP, immunoprecipitated. GAPDH, loading control.

### UBA and UIM domains of USP28 are not required for p53 activation

The N-terminus of USP28 is not essential for its interaction with 53BP1, but it contains two distinct regions: the Ubiquitin-associated domain (UBA) and the Ubiquitin-interacting motif (UIM) (**Fig. 3B**). These domains are proposed to bind ubiquitinated proteins and may play a role in substrate recognition of USP’s ^30^. To evaluate whether UBA and UIM domains are necessary for p53 stabilization under mitotic stress, we isolated two single clones of truncation mutants lacking UBA (92–1045) or both UBA and UIM (157–1045) (**Fig. 3B, S5)**. All clones, including those expressing full-length USP28 and the truncated variants, showed comparable expression levels. Unexpectedly, all tested mutants were able to stabilize p53 and induce p21 expression, similar to wildtype RPE1 cells. These findings suggest that UBA and UIM-mediated substrate recognition by USP28 is dispensable for p53 stabilization in response to mitotic stress.

### Cancer-associated mutations in USP28 impair mitotic stress response

USP28 is frequently mutated in cancer and are classified as tumor suppressor (**Fig. 5A**). Frameshift mutations in USP28 disrupt protein expression and render cancer cells insensitive to mitotic stress ^13^. While frameshift mutations are known to impair USP28 function, most cancer-associated mutations in USP28 are missense mutations, substituting a single amino acid (**Fig. 5A, B**). We hypothesized that missense mutations in USP28 may desensitize cancer cells to mitotic stress, contributing to genome instability, which is a hallmark of cancer.

**Figure 5:**
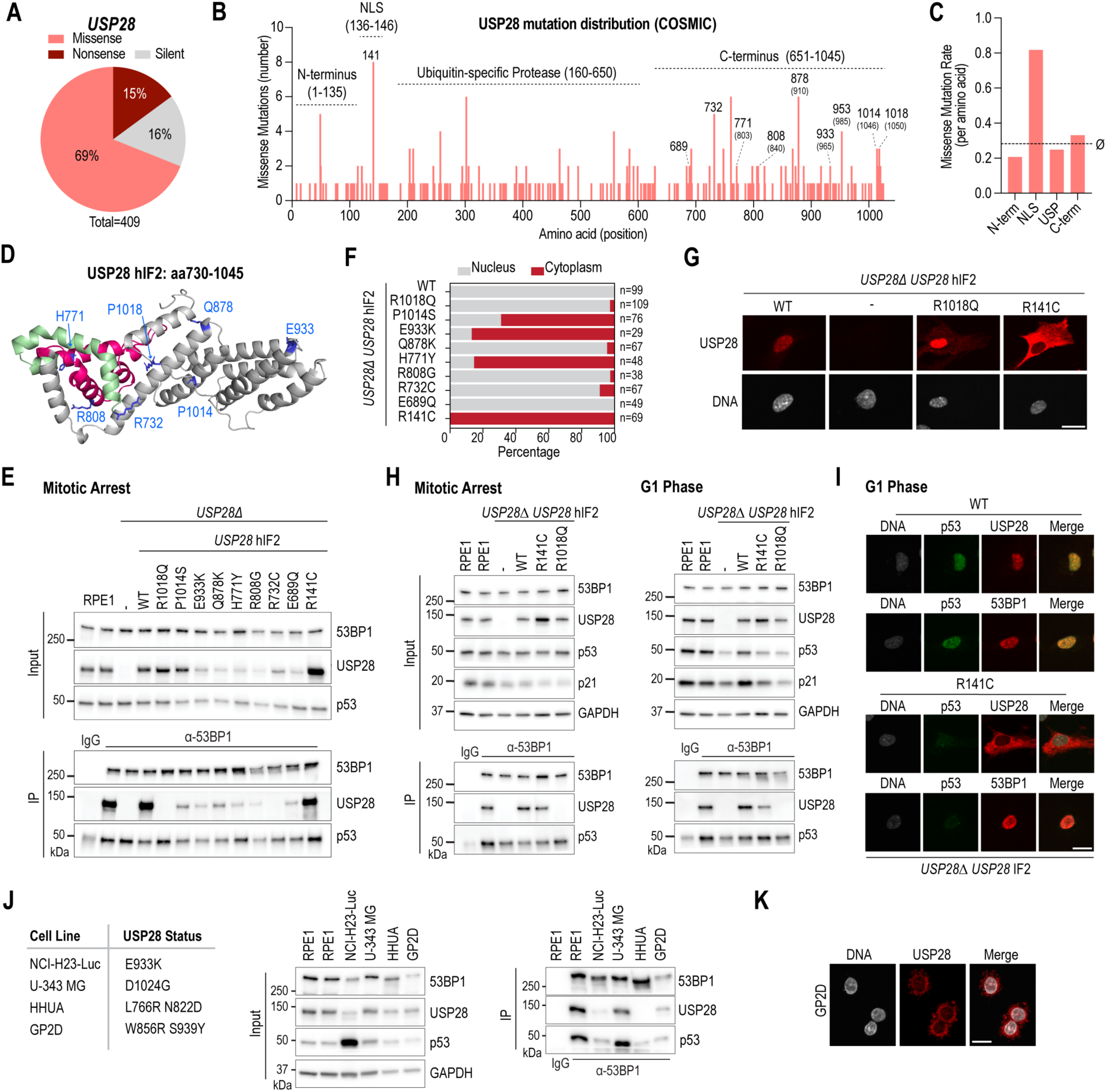
Cancer-associated mutations in USP28 impair mitotic stress response (A) Pie chart summarizing USP28 point mutations identified in tumors (COSMIC database). (B) Distribution of missense mutations across USP28 regions. Numbers indicate amino acid positions in USP28^hIF2^; corresponding USP28^hIF1^ positions are in brackets. Regions quantified in (C) are highlighted. **(C)** Missense mutation rate per amino acid in distinct domains of the USP28 protein. The dotted line shows the average (ø) across the entire gene. **(D)** AlphaFold-predicted structural model of the USP28 C-terminus highlighting residues frequently mutated in cancer. **(E)** Immunoprecipitation of 53BP1 from mitotically arrested cells to evaluate complex formation with USP28 and p53. *USP28*^hIF2^ variants with cancer-associated mutations were expressed in *USP28Δ* RPE1 cells. Unmodified RPE1 cells were used as a control. Inputs are soluble supernatants. IP, immunoprecipitate. GAPDH, loading control. **(F)** Quantification of nuclear vs. cytoplasmic localization for cancer-associated USP28 mutants (highlighted in B, D). **(G)** Representative images of wildtype and mutant USP28 localization. Scale bar: 10 µm. **(H)** Immunoprecipitation analysis of 53BP1 and p53 in mitotically arrested and post-mitotic cells expressing wildtype or mutant *USP28*^hIF2^ (R141C, R1018Q). Experimental conditions as described in Figure 1F. Inputs are soluble supernatants. IP, immunoprecipitate. GAPDH, loading control. **(I)** Immunostaining of USP28 and 53BP1 following release from mitotic arrest into G1 phase. Conditions for identifying G1 phase cells are detailed in Figure 1F. The localization of the wildtype and R141C mutant of USP28 is compared with 53BP1. Scale bar: 10 µm. **(J)** Immunoprecipitation analysis of 53BP1 in four cancer-derived cell lines with C-terminal USP28 mutations. Inputs are soluble supernatants. IP, immunoprecipitate. GAPDH served as a loading control. **(K)** Immunostaining of GP2D cells to assess subcellular localization of endogenous USP28. Scale bar: 10 µm.

To test this hypothesis, we first analyzed the frequency distribution of missense mutations across the USP28 coding region. We found that amino acids in the C-terminus (aa651-1045) were more frequently mutated than those in the USP domain (aa160-650) or the N-terminus (aa1-135) (**Fig. 5C**). Notably, a single amino acid within the NLS (aa135-145), R141C, was the most mutated, suggesting that nuclear localization is critical for USP28 function.

To assess the impact of the most frequent C-terminal mutations and the NLS mutation, we expressed mutated USP28^hIF2^ transgenes in *USP28Δ* RPE1 cells (**Fig. 5B, D**). We observed three distinct phenotypes. First, mutations between amino acids 689 and 933 led to protein destabilization, as evidenced by reduced expression levels (**Fig. 5E**), proposing that the reduced expression could desensitize to prolonged mitosis. Second, the R141C mutation within the NLS caused aberrant cytoplasmic localization (**Fig. 5F, G**). Intriguingly, several mutations in the C-terminus also resulted in exclusion from the nucleus, consistent with the cytoplasmic localization of previously described mutants (G903L, P953L) (**Fig. 2D, 5F)**. Proline 953 is also often mutated in cancer (**Fig. 5B),** supporting the possibility that USP28 is excluded from the nucleus in some cancers. Third, the R732C and R1018Q mutants failed to interact with 53BP1 regardless of their nuclear localization (**Fig. 5E-G**), which further supports the possibility that cancer impairs mitotic stress response by interfering with mitotic stopwatch complex formation and function.

To investigate the functional consequences of these mutations, we examined p53 stabilization following prolonged mitosis. We arrested R141C (nuclear exclusion) and R1018Q (failure to interact with 53BP1) mutants in mitosis and subsequently released them into the next cell cycle (**Fig. 5H**). Both mutants failed to stabilize p53 in G1 phase. The inability of the R1018Q mutant to activate p53 is likely due to the lack of interaction with 53BP1. While the R141C mutant formed a stable complex with 53BP1 and p53 during mitotic arrest (**Fig. 5E, H**), it failed to stabilize p53 after mitotic exit, which might be caused by the spatial separation of cytoplasmic USP28 and nuclear 53BP1 (**Fig. 5I**).

To confirm that the tested mutations have similar defects in cancer, we identified four cancer-derived cell lines that have either one homogenous or two heterogenous missense mutation in the C-terminus of USP28 (**Fig. 5J**). We found that three cell lines had reduced level of USP28 expression, which is in line with our finding from transgenic RPE1 cells (**Fig. 5E**). Furthermore, we found that USP28 either failed to interact with 53BP1 (HHUA) or had a lower degree of interaction (NCI-H23) (**Fig. 5J**). Immunostaining revealed that USP28 failed to localize in the nucleus in GP2D cells (**Fig. 5K**).

In conclusion, all tested cancer-associated missense mutations in the C-terminus of USP28 resulted in nuclear exclusion, failure to interact with 53BP1, or protein destabilization; phenotypes which we also observed in cancer-derived cell lines. These mutations prevent the activation of p53 following extended mitosis, highlighting their role in disrupting the mitotic stress response in cancer cells.

### Destabilization of USP28 reduces sensitivity to mitotic stress

To assess if the reduced USP28^hIF2^ amount in cancer cells alters the sensitivity to prolonged mitosis, we generated single-cell clones with homogenous but different expression levels of USP28^hIF2^ (**Fig. 6A, B**). We selected three clones: Clone 1 (#1), which exhibited slightly higher USP28^hIF2^ expression than wildtype RPE1 cells; Clone 2 (#2), with expression levels similar to wildtype; and Clone 3 (#3), which expressed lower levels of USP28^hIF2^ (**Fig. 6B-D**). Following a 16-hour mitotic arrest, all three clones demonstrated complex formation between USP28^hIF2^, 53BP1, and p53, though the abundance of USP28^hIF2^ within these complexes correlated with the level of USP28^hIF2^ expression (**Fig. 6D**).

**Figure 6:**
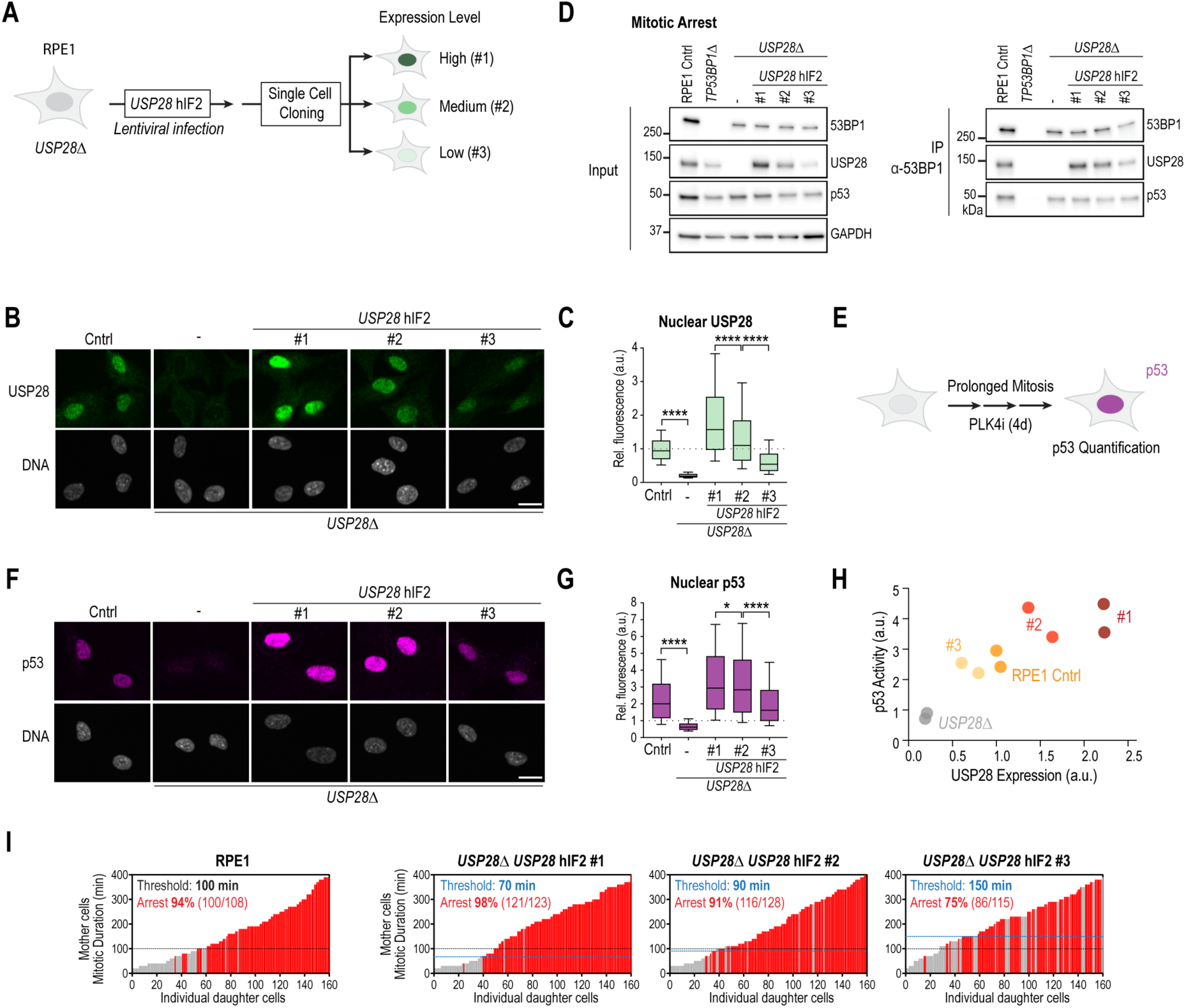
USP28 confers concentration-dependent sensitivity to prolonged mitosis **(A)** Schematic of single-cell clone isolation expressing the *USP28*^hIF2^ transgene. Clones #1, #2, and #3 show homogenous expression. **(B)** Immunostaining of RPE1 control, *USP28Δ*, and *USP28*^hIF2^-expressing clones. Clones were isolated as in (A). Scale bar: 10 µm. **(C)** Quantification of USP28 staining from (B). Box-and-whisker plots show mean and 10–90 percentile (error bars), normalized to RPE1 levels (dotted line at 1). One-way ANOVA: **P < 0.01; ***P < 0.001; ****P < 0.0001. n > 2,500 cells/condition. **(D)** Immunoprecipitation of 53BP1 from mitotically arrested cells to assess complex formation with USP28 and p53. Clones expressing different *USP28*^hIF2^ levels were analyzed. Unmodified and *TP53BP1Δ* RPE1 cells were used as controls. Inputs are soluble supernatants. IP, immunoprecipitate. GAPDH, loading control. **(E)** Schematic of assay to assess sensitivity to prolonged mitosis. Cells were treated with PLK4 inhibitor for 3 days and stained for p53 activation. **(F)** Immunostaining of RPE1 control, *USP28Δ*, and *USP28*^hIF2^ clones for p53 following prolonged mitosis. Scale bar: 10 µm. **(G)** Quantification of p53 staining from (F). Data shown as box-and-whisker plots, normalized to untreated RPE1 (dotted line at 1). One-way ANOVA: *P < 0.05; **P < 0.01; ***P < 0.001; ****P < 0.0001. n > 2,500 cells/condition. **(H)** Correlation plot showing USP28 levels (from C) versus p53 stabilization (from G) after prolonged mitosis. Each point represents mean of replicates. See also Figure S6D. **(I)** Imaging-based assay (as described in Figure 1E) used to examine how USP28 expression affects the mitotic duration threshold required to induce cell cycle arrest. RPE1 control graph reused from Figure 1E.

To determine whether the 53BP1-USP28^hIF2^ complexes could activate p53 in response to prolonged mitosis, we treated cells with a PLK4 inhibitor (PLK4i) to induce centrosome depletion and thereby prolong mitosis (**Fig. 6E**) ^9,33^. PLK4 inhibition does not affect the first mitosis. The second and following mitoses are moderately prolonged (50–120 minutes) ^13^. Since RPE1 cells complete a cell cycle in approximately 20 hours, they typically experience one or two moderately prolonged mitoses within three days of treatment. After three or four days of PLK4i treatment, we analyzed p53 stability through immunostaining (**Fig. 6F, G; S6A-C)** and observed that p53 activation levels correlated with USP28^hIF2^ expression levels (**Fig. 6H; S6D)**.

To further investigate whether the sensitivity to mitotic stress depends on USP28 expression levels, we tracked individual cells by live cell imaging and assessed the response to single prolonged mitosis (**Fig. 1E**). Clone 1, with approximately 1.5-fold higher USP28^hIF2^ expression than wildtype cells, showed increased sensitivity to mitotic arrest, while Clone 3, with roughly half the wildtype USP28^hIF2^ level, exhibited reduced sensitivity (**Fig. 6I**). Specifically, the threshold for inducing cell cycle arrest in response to prolonged mitosis was approximately 100 minutes in wildtype cells, 70 minutes in Clone 1, 80 minutes in Clone 2, and 150 minutes in Clone 3. These findings indicate that cellular sensitivity to mitotic stress is modulated by the expression level and stability of USP28^hIF2^, establishing USP28 as a limiting factor for mitotic stress response.

## Discussion

USP28 has been described as a tumor suppressor and oncogene ^11,34^. Here, we demonstrate that in both untransformed and cancerous p53-wildtype cells, USP28 primarily functions as a tumor suppressor by stabilizing p53 and downregulating MYC in response to mitotic stress (**Fig. 1A-D**). Unexpectedly, we found that in some cancer cell lines USP28 increases baseline p53 levels. In these cell lines, USP28 deletion not only dampens the response to mitotic stress but also weakens the response to DNA damage. Furthermore, we elucidate the molecular mechanisms by which USP28 stabilizes p53 following mitotic stress (**Fig. 2-5**) and show that these mechanisms are frequently inactivated in cancer (**Fig. 6, 7)**. These findings highlight the tumor-suppressive roles of USP28 and may have important implications for therapeutic strategies.

Using transgene expression, we found that the tumor suppressor function of USP28 is mediated by the shorter isoform 2 of USP28, but not the longer isoform 1, which is frequently considered as canonical isoform (**Fig. 1G-J**). The two isoforms differ by 32 amino acids that are expressed from exon 19. AlphaFold modeling predicted that this region forms an alpha helix (**Fig. 1I**). A possible explanation is that the additional 32 amino acids of the longer isoform 1 disrupt the interaction surface between the C-terminus of USP28 and 53BP1. In line with this model, we identified several point mutations (G903R and P953L) and truncation mutations in the same C-terminal region of USP28 (D1003fs, C-Δ13) that impair the interaction with 53BP1 (**Fig. 2F; 5E, J)** and consequently fail to stabilize p53 in response to mitotic stress (**Fig. 5H**). Surprisingly, isoform 2 of USP28 is more dominantly expressed than isoform 1 across 29 cancer cell lines from 10 different tissues (**Fig. 1K, L; S2E, G, H)**. Additionally, we observed a similar expression pattern in various mouse organs, including liver, kidney, heart, muscle, and lung (**Fig. 1M; S2F)**. The brain is the only organ identified in mice in which the longer isoform or other Exon 19 expressing isoforms were predominantly expressed. These findings suggest that the shorter isoform (*USP28*^hIF2^; *USP28*^mIF1^) is the canonical isoform, while the longer isoform (*USP28*^hIF1^; *USP28*^mIF2^) is context-specific. Further studies are needed to explore the precise function of the longer isoform (*USP28*^hIF1^; *USP28*^mIF2^) and its expression in different cell types.

We further investigated the minimal region required for the interaction between USP28 and 53BP1. Surprisingly, while the C-terminus of USP28 is required for binding to 53BP1, it is not sufficient on its own (**Fig. 3B, D**). The minimal USP28 fragment that was able to interact with 53BP1 contained the USP domain but lacked the N-terminal domain, which includes ubiquitin-binding motifs UBA and UIM and an NLS (**Fig. 3D**). The USP domain of USP28 is interrupted by a dimerization arm-specific to USP25 and USP28 (**Fig. 3E**) ^28,29^. Disruption of dimerization through two mutations (V541E, L545E) impaired USP28’s interaction with 53BP1, suggesting that USP28 must form a dimer to interact with 53BP1 (**Fig. 3B, G**). These results suggest two possible mechanisms: either dimerization of USP28 is required for the interaction between the C-terminus of USP28 and 53BP1, or the dimerization arms create a second binding surface for 53BP1. Using inducible dimerization domains (DmrB), we were able to replace the function of the dimerization arm, showing that dimerization of the C-terminus of USP28 is sufficient for 53BP1 binding (**Fig. 3H-J; S4F-I)**. Furthermore, we found that the interaction of the USP28 C-terminus and 53BP1 is PLK1-dependent and critical for the transfer of mitotic stress signaling to the progenitor cells (**Fig. 4**). Our findings raise new questions about the stoichiometry of the mitotic stopwatch complex. Previous studies have shown that 53BP1 forms oligomers ^35,36^ and p53 exists as a tetramer ^37^, which implies that the mitotic stopwatch complex may have a higher structural complexity.

USP28 is frequently impaired in cancer (**Fig. 5A**). Cancer-associated missense mutations in USP28 prevent p53 stabilization following mitotic stress. We identified five functional domains of USP28 essential for mitotic stress response: the USP domain, 53BP1 interaction, dimerization, protein stability, and nuclear localization (**Fig. 7A**). Our work revealed that defects in three of these domains are associated with frequent missense mutations in cancer (**Fig. 7B**). We found that the two out of ten (20%) tested cancer-associated mutations (R732C and R1018Q) impair the interaction with 53BP1 and fail to stabilize p53 following mitotic stress, highlighting the importance of this interaction (**Fig. 5E, H**). Several cancer-associated mutations in the C-terminus also result in reduced USP28 stability, which likely dampens the mitotic stress response in cancer cells (**Fig. 5E, 6)**. The most frequent mutation in USP28 occurs within the NLS domain (**Fig. 5B**), impairing nuclear localization and the mitotic stress response (**Fig. 5F-H**). While nuclear exclusion does not prevent mitotic stopwatch complex formation, it impairs p53 stabilization in G1 phase, possibly due to the separation of USP28 in the cytoplasm and 53BP1 in the nucleus (**Fig. 5H, I**). Interestingly, mutations in the C-terminus also cause nuclear exclusion, but the underlying mechanism remains unclear, as no NLS sequence is detected in this region.

**Figure 7:**
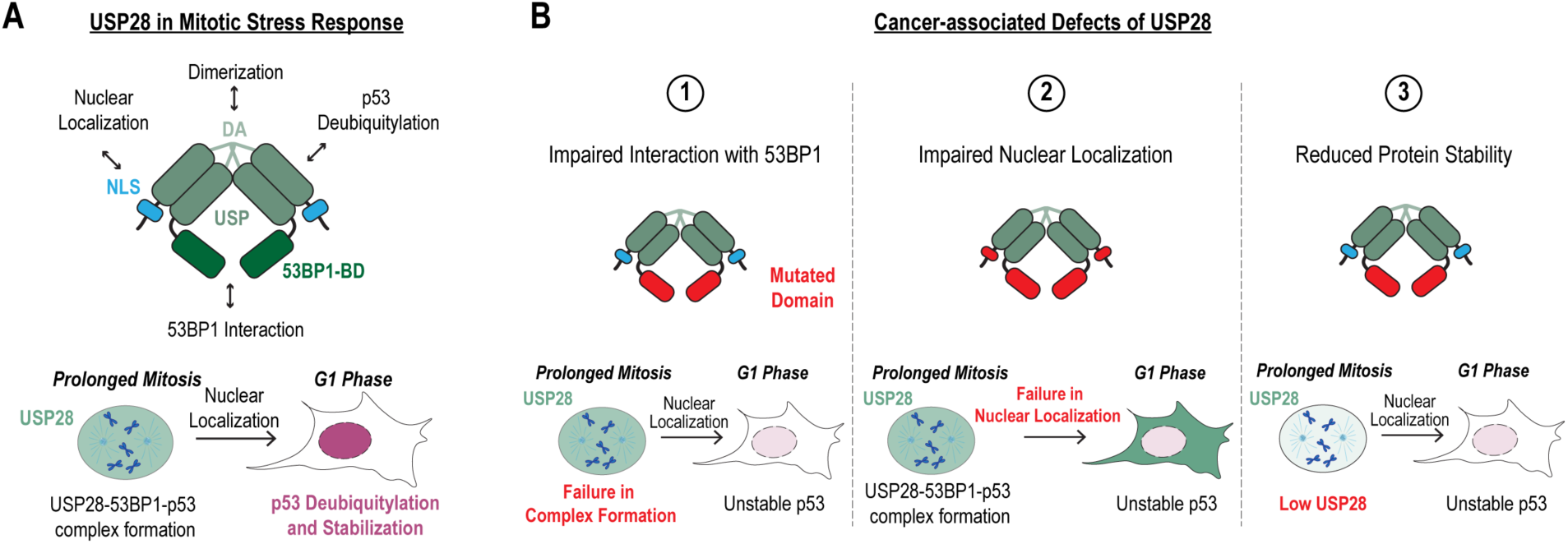
Tumor-suppressive mechanisms of USP28 **(A)** Model illustrating USP28 dimer structure. Essential functions that are required for prolonged mitosis-mediated p53 stabilization are highlighted. DA, Dimerization arm; USP, Ubiquitin-specific protease; 53BP1-BD, 53BP1-binding domain; NLS, Nuclear localization sequence. **(B)** Schematic summarizing cancer-associated functional impairments caused by mutations in USP28: 1. Impaired interaction with 53BP1, which are caused by mutations in the 53BP1-BD domain. 2. Impaired nuclear localization, which is caused by mutations in the NLS or 53BP1-BD domain. 3. Reduced USP28 stability, which is caused by mutations in the 53BP1-BD domain.

Several studies have identified USP28 as an oncogene in specific cancer contexts ^1,2,4,38,39^. A recent study provided mechanistic insight into this function, showing that DNA damage activates ATM, which triggers the disassembly of USP28 dimers into monomers ^3^. These monomeric forms then stabilize MYC, promoting oncogenic proliferation³. In contrast, our findings reveal a distinct pathway in which mitotic stress enhances the interaction between USP28 and 53BP1, leading to the stabilization of p53 and the downregulation of MYC. We further observed that cancer cells frequently disrupt the USP28–53BP1 interaction, potentially as a strategy to bypass p53-mediated cell cycle arrest or apoptosis (**Fig. 5**). Notably, USP28 mutants found in cancer may retain the ability to stabilize MYC, thus maintaining oncogenic signaling. Consistent with this, mutations are more frequently found in the C-terminal region of USP28— which mediates 53BP1 binding—than in the catalytic USP domain required for MYC stabilization (**Fig. 5C**). Given that USP28-mediated activation of p53 is critical for mounting an effective response to mitotic stress response and in some cancers to DNA damage response, disruption of its interaction with 53BP1 could contribute to therapeutic resistance, particularly against anti-mitotic and DNA-damaging agents.

A wide spectrum of cellular defects and environmental insults—including mitotic machinery dysfunction, DNA replication stress, aneuploidy, heat and osmotic stress, oncogene activation, irradiation, increased cell size, and viral infections—can lead to mitotic errors or DNA damage, ultimately promoting genomic instability ^14,40–52^. We propose that USP28 acts as a central node in the cellular stress response network, particularly in the context of mitotic and DNA damage stress. By fine-tuning the cellular threshold for responding to both intrinsic and extrinsic stressors, USP28 may play a key role in maintaining genomic stability and tissue integrity.

## Materials and Methods

### Chemical inhibitors

The chemical inhibitors and concentrations used in this study were: Centrinone (PLK4i; LCR-263; 150 nM; MedChem Express); Nocodazole (0.1 µg/ml; Sigma-Aldrich); Monastrol (100 µM; Tocris Bioscience); B/B Homodimerizer (100 nM; Takara); Doxorubicin (1-1000 nM; Cell Signaling).

### Antibodies

The following antibodies were purchased from commercial sources, with their working concentrations indicated in parentheses: anti-53BP1 (1:5000; Novus Biologiclas Cat# NB100-304, RRID:AB_10003037), USP28 (1:1000; Abcam Cat# ab126604, RRID:AB_11127442), USP28 (1:100; Sigma-Aldrich Cat# HPA006778, RRID: AB_1080520), p53 (1:1000; Santa Cruz Biotechnology Cat# sc-126, RRID:AB_628082), p21 (1:1000; Cell Signaling Technology Cat# 2947, RRID:AB_823586), anti-GAPDH (1:1000; Cell Signaling Technology Cat# 5174, RRID:AB_10622025), anti-Myc 9E10 (1:1000; Sigma-Aldrich Cat# M4439, RRID: AB_439694), anti-Histone H3.3 (1:1000; Abcam Cat#ab5176, RRID: AB_304763); anti-Flag M2 (1:1000; Sigma-Aldrich Cat#F1804-200UG, RRID: AB_262044); Anti-phospho-Histone H2A.X (1:2000; Millipore Cat# 05-636, RRID:AB_309864); IgG Rabbit (1:5000; Vector Laboratories Cat# I-1000-5). Secondary antibodies were purchased from Jackson ImmunoResearch and GE Healthcare.

### Cell lines

All cell lines used in this study are described in **Table S1**. RPE1 (hTERT RPE-1), MCF7, CHP212, G401, A375, HCT116, RKO, U20s, SJSA1, 769P, H460, SH-SY5Y, N2A and MP41 were obtained from the American Type Culture Collection (ATCC); CHP134 and Mel-202 were obtained from Sigma-Aldrich (ECACC general collection) and G402, A549, LU99, Caki-1, ONDA9, A172 and OVTOKO cell line from Japanese Collection of Research Biosources (JCRB) Cell Bank. LOX-IMVI was obtained from the NCI-60 collection. Cell lines were cultured in the recommended growth medium at 37°C and 5% CO_2_, supplementing the growth media with 100 IU/ml penicillin and 100 µg/ml streptomycin. To inhibit cells in mitosis, cells were treated with either 100 ng/ml of Nocodazole (Sigma-Aldrich) or 100 µM Monastrol (Tocris Bioscience) for the indicated amounts of time.

The RPE1 *USP28Δ* cell line has been described ^13^. The following transgenes were stably integrated into the genome of *USP28Δ* RPE1 cells using lentivirus constructs (**see Table S2**): H2B-mRFP (EF1alpha promoter); USP28 mutant transgenes (UbC promoter). Viral particles were generated by transfecting the lentiviral plasmid (**see Table S2**) into HEK-293T cells using Lenti-X Packaging Single Shots (Clontech, Cat# 631276). 48 hours after transfection, virus-containing culture supernatant was harvested and added to the growth medium of cells in combination with 8 µg/ml polybrene (EMD Millipore). Populations of each cell line were selected by antibiotics (Neomycin, 400 µg/ml) or FACS. Single clones were isolated in 96-well plates, and single colonies with USP28 transgenes were analyzed using immunofluorescence assays. Genomic DNA from the transgene clones was isolated using the Quick-DNA Microprep Kit (ZYMO RESEARCH, cat# D3021), and the USP28 transgenes were then sequenced to identify mutations.

### Plasmid construction

All plasmids used in this study are described in **Table S2**. USP28 mRNA Isoform-2 (NCBI reference sequence: NM_001346258.2), which lacks Exon 19 compared to the canonical isoform-1 mRNA sequence (NCBI reference sequence: NM_020886.4) were cloned in lentiviral expression vectors. We optimized the UbC promoter length (deletion of 245 bp from the N-terminal region of the promoter) to reduce USP28 expression. This plasmid served as the backbone for generating USP28 mutant transgene constructs using site-directed mutagenesis with Phusion® High-Fidelity DNA Polymerase (NEB Bio-lab, Cat#M0530L). Cluster mutants were synthesized (gBlocks, IDT) and cloned into the vector.

### Immunofluorescence

For immunofluorescence, approximately 5,000 cells per well were seeded into 96-well imaging plates (SCREENSTAR, cat# 655866) one day before fixation. Cells were fixed in 100µl ice-cold methanol for 7 minutes at −20°C. Cells were washed twice with washing buffer (PBS containing 0.1% Triton X-100) and blocked with blocking buffer (PBS containing 2% BSA, 0.1% Triton X-100 and 0.1% sodium azide) for 2 hours at 37°C or overnight at 4°C. After blocking, cells were incubated for 1-2 hours with indicated primary antibody in fresh blocking buffer (concentrations as indicated above). Cells were washed three times with washing buffer, prior to 1-hour incubation with the secondary antibody and DNA staining with Hoechst 33342 dye. Finally, cells were washed three times with washing buffer. Images were acquired on a CellVoyager CQ1 spinning disk confocal system (Yokogawa Electric Corporation) equipped with a 40X (0.95 NA) objective and a 2000×2000 pixel sCMOS camera (ORCA-Flash4.0V3, Hamamatsu Photonics). Image acquisition was performed using CQ1 software.

### USP28 transgene activity

To test the activity of wildtype and mutant versions of USP28, transgene-expressing *USP28Δ* RPE1 cells were treated for 3 or 4 days with the PLK4 inhibitor Centrinone (150 nM) or DMSO as control ^33^. On the second or third day of treatment approximately 7,000 cells were plated into a 96-well imaging plate while maintaining Centrinone or DMSO treatment. Cells were fixed on day 3 or 4 and stained with Hoechst DNA dye and antibodies for USP28 and p53. Images were acquired on a CQ1 spinning disk confocal system (Yokogawa Electric Corporation). Image acquisition was performed using CQ1 software from Yokogawa as previously described. Signal intensities were quantified using Yokogawa Pathfinder software.

### Live cell imaging

Live cell imaging was performed on the CellVoyager CQ1 spinning disk confocal system (Yokogawa Electric Corporation) equipped with a 40X 0.95 NA U-PlanApo objective and a 2000×2000 pixel sCMOS camera (ORCA-Flash4.0V3, Hamamatsu Photonics) at 37°C and 5% CO_2_. Image acquisition and data analysis were performed using CQ1 software and ImageJ, respectively.

### Mitotic stopwatch assay

All imaged cell lines were engineered to express H2B-RFP (see **Table S1**). One day before imaging, cells were seeded into 96-well SCREENSTAR (Greiner, cat# 655866) plates at 2,000-4,000 cells/well. On the day of the experiment, asynchronous cells were first imaged for 4-6h in 100 µM Monastrol. Images of H2B-RFP were acquired with 5 x 2μm z-sections in the RFP channel (25% power, 150 ms) at 10-minute intervals. During Monastrol treatment, cells enter at different times into mitosis and delay in prometaphase. This generates a set of mother cells that experience different mitotic durations prior to Monastrol washout. After Monastrol washout, the mother cells completion of mitosis was imaged at 10-minute intervals for 2 hours and the fate of the resulting daughter cells was imaged at 20-minute intervals; daughter was tracked for 48-72 hours. The fate of the daughter cells was classified into ‘arrest’, ‘death’ or ‘proliferate’. The threshold for mitotic stopwatch activation was determined as the mitotic duration for which more than 50% of the daughter cells were arrested.

### Competition Assays

Wildtype cells and a pool of heterogenous *USP28* or *TP53BP1* knockout cells were mixed and seeded into 10 cm plates at 100,000-300,000 cells/plate and treated with PLK4i (150 nM), Doxorubicin (10 nM) or DMSO as a control. Cells were maintained for 8 days and passaged to avoid complete confluence. After 8 days cells were harvested, and genomic DNA purified. The genomic region targeted in *USP28* or *TP53BP1* was sequenced, and the knockout ratio was calculated by the indel decomposition software TIDE (http://shinyapps.datacurators.nl/tide/). The ratios PLK4i/DMSO and Doxorubicin/DMSO were calculated and blotted for each cell line.

### Proliferation assays

For proliferation analysis, cells were seeded into 6 well plates in triplicate at 25,000 cells/well and treated with the indicated inhibitors or DMSO as a control. At 96-hour intervals, cells were harvested, counted and, for passaging assays, re-plated at 25,000 cells/well. Cell counting was performed using a TC20 automated cell counter (Bio-Rad).

### MYC Stability assay

For the MYC stability analysis, 1X10^5^ cells were plated into a 6-well plate. After 24 hours, cycloheximide was added to the cell culture medium at a concentration of 100 μg/ml. Cells were then harvested at indicated time points. Following this, the cells were lysed, and 5 µg of the lysate was immunoblotted using the specified antibodies.

### Artificial homodimerization assay of USP28

Single DmrB, an inducible homodimerization domain (Takara), was amplified by PCR and cloned together with a FLAG tag into USP28 expressing lentiviral constructs upstream of the gene fragment USP28_aa580-1045 and USP28_aa651-1045 (*UbC^pro^-3xFLAG-DmrB-USP28-(aa580-1045); UbC^pro^-3xFLAG-DmrB-USP28-(aa651-1045)).* The transgenes were stably integrated into the *USP28Δ* RPE1 cell line genome using lentiviral constructs. To induce USP28 homodimerization, cells were treated with 100 nM of B/B Homodimerizer along with Nocodazole (100 ng/ml) for 16 hours. After collecting the treated mitotic cells, they were subjected to immunoprecipitation.

### Immunoblotting

For immunoblotting, cells were cultured in 15 cm plates, harvested at 80% confluence, and lysed by sonication in Lysis buffer (20 mM Tris-HCl (pH 7.5), 50 mM NaCl, 0.5% Triton X-100, 5 mM EGTA, 1 mM dithiothreitol, 2mM MgCl2) plus protease and phosphatase inhibitor cocktail (Thermo Fisher Scientific). After 15 minutes of centrifugation at 15,000 x g and 4°C, the cell extract was separated from the cell debris and stored at −80°C until use. Extract concentrations were measured based on protein standard concentration (Bio-Rad Protein Assay) and 5-10 μg protein was loaded per lane on Mini-PROTEAN gels (Bio-Rad) and transferred to PVDF membranes using a TransBlot Turbo system (Bio-Rad). Blocking and antibody incubations were performed in TBS-T + 5% non-fat dry milk. Detection was performed using HRP-conjugated secondary antibodies (GE Healthcare) with SuperSignal West Femto (Thermo Fisher Scientific) substrates. Membranes were imaged on a ChemiDoc MP system (Bio-Rad).

### Immunoprecipitation

For immunoprecipitation assays cells were arrested in mitosis by treatment with 100 ng/ml (0.33 or 0.66 µM) Nocodazole for 8 or 16 hours. 1-2 million cells were harvested and washed with PBS. Cells were re-suspended in lysis buffer (20 mM Tris/HCl pH 7.5, 50 mM NaCl, 0.5% Triton X-100, 5 mM EGTA, 1 mM dithiothreitol, 2 mM MgCl_2_ and EDTA-free protease inhibitor cocktail (Roche)) and lysed in an ice-cold sonicating water bath for 5 minutes. After 15-minute centrifugation at 15,000 x g and 4 °C, soluble lysates were collected, and protein concentrations were quantified. Equal amounts of lysates (1-2 mg) were incubated with anti-53BP1(Novus Biologiclas, Cat# NB100-304, RRID:AB_10003037) for 2 hours at 4°C and subsequently with Protein A magnetic bead (Thermo Fisher Scientific, Cat# 88845) for 1 hour at 4 °C. The beads were washed five times with lysis buffer and re-suspended in SDS sample buffer. For immunoblotting, equal volumes of samples were run on Mini-PROTEAN gels (Bio-Rad) and transferred to PVDF membranes using a TransBlot Turbo system (Bio-Rad). Blocking and antibody incubations were performed in TBS-T plus 5% nonfat dry milk. Immunoblotting was performed as described above.

For immunoprecipitation assays in G1 phase following prolonged mitosis, cells were first arrested in mitosis with 100 ng/ml Nocodazole for 8 or 16 hours. To allow cells to exit mitosis and enter G1 phase, mitotic cells were washed four times with PBS and replated onto 15 cm dishes. Plated cells were allowed to exit mitosis and enter G1 phase. After 6 hours G1 phase cells were harvested, and immunoprecipitation assays were performed as described above.

### RT-PCR Analysis

Total RNA was prepared by lysing 1X10^7^ cells and 100 mg of mouse tissues with RNeasy Plus Kit (Qiagen) and TRIzol (Thermo Fisher Scientific) respectively. Then, a total of 500 ng of RNA was used to generate cDNA by using a PrimeScript™ II 1st strand cDNA Synthesis Kit (TAKARA BIO, Japan, 6210A) with oligo dT primer. Subsequently, 1 µl of 1st strand cDNA was used as a template for PCR analysis. Electrophoresis was performed after PCR to confirm the amplification of a PCR product from the designed primer set. The primer sequences are listed in **Table S3**.

### Quantification and Statistical Analysis

All statistical analyses were performed using GraphPad Prism (version 10.4.1). For comparisons between more than two groups, one-way analysis of variance (ANOVA) was used, followed by Dunnett’s post-hoc tests where specified in figure legends. In Figures 2D, 6C, 6G and S6C data are presented as box-and-whisker plots showing the mean and 10–90 percentiles. One-way ANOVA was used to determine statistical significance between groups, with p-values indicated as follows: P < 0.05 (*), P < 0.01 (**), P < 0.001 (***), and P < 0.0001 (****). For large-scale single-cell quantifications (e.g., >1,000 or >2,500 cells/condition), individual cells were treated as independent observations. Normal distribution was assumed given large sample sizes. Replicate means were used for correlation analysis shown in Figure 6H and S6D. Sample sizes are indicated in the figures or corresponding figure legends.

For binary imaging-based classification of cell fates (e.g., arrest vs. division in Figures 1E, 1J, 2C, 6I), categorical distributions were plotted, and thresholds (such as mitotic duration cutoffs) were defined based on the point at which >50% of daughter cells exhibited arrest. Immunoblot quantifications were semi-quantitatively evaluated, with fold changes relative to DMSO-treated controls noted directly in the figures (e.g., >1.5× for p53/p21 induction, <0.5× for MYC repression).

## Materials Availability Statement

The materials (cell lines, plasmids) generated for this article will be shared on reasonable request to the corresponding authors.

## Acknowledgment

We thank Tadashi Yamamoto and Tomomi Kiyomitsu for sharing reagents; the OIST Animal Resource Section for support; Muhammad Hamzah for feedback on the manuscript and members of the Cell Proliferation and Gene Editing Unit for discussion. This work was supported by the Okinawa Institute of Science and Technology and the Japan Society for the Promotion of Science [KAKENHI, 23K05773 to F.M. and 24K09461 to M.O.].

## Author contribution

Conceptualization: H.B., M.O., F.M.; Funding acquisition: F.M., M.O.; Investigation: H.B., F.M.; Methodology: H.B., E.F.Y.N., M.O., F.M.; Resources: F.M.; Writing – original draft: H.B., F.M.; Writing – review & editing: H.B., E.F.Y.N., M.O., F.M.

