## Supplemental Figures for "Cancer-Associated USP28 Missense Mutations Disrupt 53BP1 Interaction and p53 Stabilization"

**Affiliations:**
[1]Okinawa Institute of Science and Technology Graduate University, Okinawa 904-0495, Japan.

**Running Head:** USP28 in mitotic stress response

Figure S1

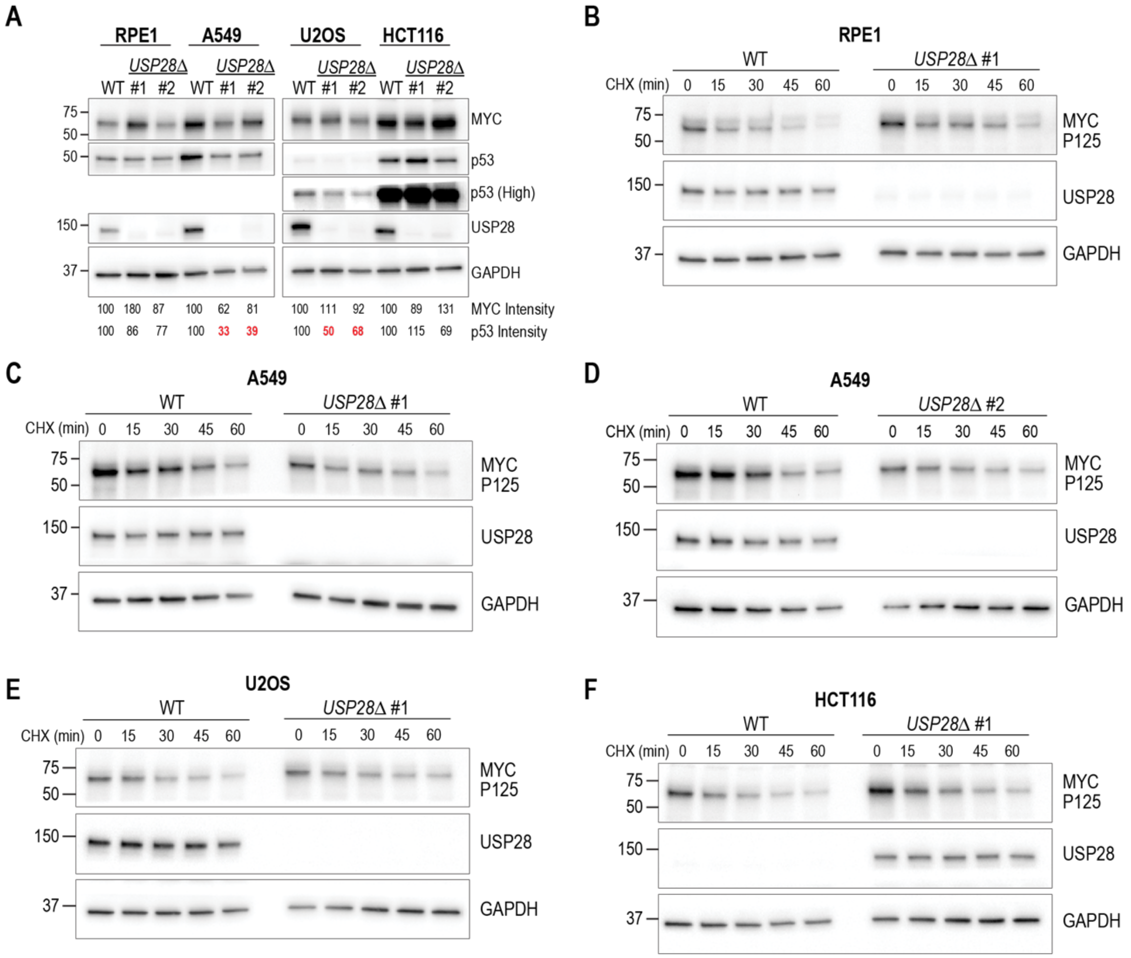


**Figure S1: Effect of USP28 deletion on p53 and MYC stability.**
**(A)** Analysis of p53 and MYC expression in cell lines with the indicated geneotypes. GAPDH served as loading control.
**(B-F)** Analysis of MYC stability in control and USP28 deleted clones following cycloheximide treatment in RPE1 (B), A549 (C, D), U2OS (E) and HCT116 (F) cells. GAPDH served as loading control.

### Figure S2

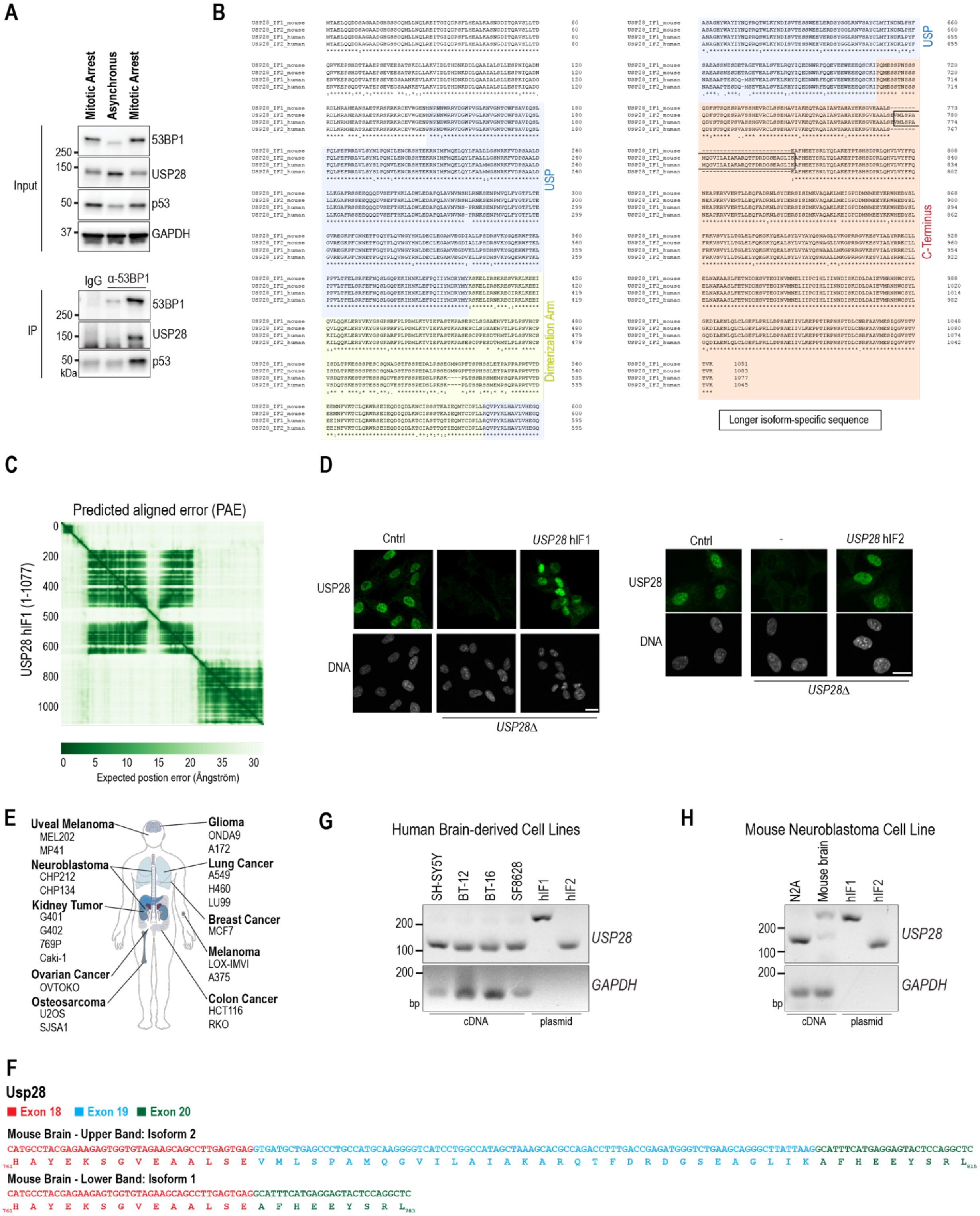
A
Mitotic Arrest
Asynchronous
Mitotic Arrest
Input
IP
53BP1
USP28
p53
GAPDH
IgG
α-53BP1
250
150
50
37
kDa
B
USP
Dimerization Arm
C-Terminus
Longer isoform-specific sequence
C
Predicted aligned error (PAE)
USP28 hIF1 (1-1077)
Expected postion error (Ångström)
D
Cntrl
USP28 hIF1
USP28 hIF2
USP28
DNA
USP28Δ
E
Uveal Melanoma
MEL202
MP41
Neuroblastoma
CHP212
CHP134
Kidney Tumor
G401
G402
769P
Caki-1
Ovarian Cancer
OVTOKO
Osteosarcoma
U2OS
SJSA1
Glioma
ONDA9
A172
Lung Cancer
A549
H460
LU99
Breast Cancer
MCF7
Melanoma
LOX-IMVI
A375
Colon Cancer
HCT116
RKO
G
Human Brain-derived Cell Lines
SH-SY5Y
BT-12
BT-16
SF8628
hIF1
hIF2
USP28
GAPDH
cDNA
plasmid
bp
H
Mouse Neuroblastoma Cell Line
N2A
Mouse brain
F
Usp28
Exon 18
Exon 19
Exon 20
Mouse Brain - Upper Band: Isoform 2
CATGCCTACGAGAAGAGTGGTGTAGAAGCAGCCTTGAGTGAGGTGATGCTGAGCCCTGCCATGCAAGGGGTCATCCTGGCCATAGCTAAAGCACGCCAGACCTTTGACCGAGATGGGTCTGAAGCAGGGCTTATTAAGGCATTTCATGAGGAGTACTCCAGGCTC
761 H A Y E K S G V E A A L S E V M L S P A M Q G V I L A I A K A R Q T F D R D G S E A G L I K A F H E E Y S R L 815
Mouse Brain - Lower Band: Isoform 1
CATGCCTACGAGAAGAGTGGTGTAGAAGCAGCCTTGAGTGAGGCATTTCATGAGGAGTACTCCAGGCTC
761 H A Y E K S G V E A A L S E A F H E E Y S R L 783

**Figure S2: Comparison of long and short isoforms of USP28 in human and mice.**
**(A)** Analysis of 53BP1 immunoprecipitates to determine complex formation with USP28 and p53 in asynchronous and mitotically arrested cells (Nocodazole, 100 ng/ml, 16 h). Inputs are soluble supernatants. IP, immunoprecipitate. GAPDH served as loading control.
**(B)** Sequence alignment of the two primary USP28 isoforms in human and mice, highlighting conserved regions. See text for more explanation.
**(C)** AlphaFold-predicted aligned error map for the full-length $USP28^{IF1}$ structure, indicating regions of higher and lower confidence in the structural model.
**(D)** Immunostaining of USP28, showing nuclear localization of endogenous USP28 and transgene-expressed $\mathit{USP28}^{hIF1}$ and $\mathit{USP28}^{hIF2}$ (clonal cell lines). The nucleus was visualized with the DNA-staining compound Hoechst 33342. Images on the right are also shown in Figure 6B. Scale bar: 10 µm.
**(E)** Overview of cancer-specific cell lines shown in Figure 1L.
**(F)** Nucleotide sequences of the expressed *USP28* mRNA confirming that isoforms with and without exon 19 are expressed in the mouse brain.
**(G-H)** RT-PCR analysis revealing that cancer-derived cell lines from human (G) and mouse (H) predominantly express the shorter isoforms $\mathit{USP28}^{hIF2}$ and $\mathit{USP28}^{mIF1}$. Plasmids containing $\mathit{USP28}^{hIF1}$ or $\mathit{USP28}^{hIF2}$ are used as control. A mouse brain sample is used for comparison (H).

Figure S3

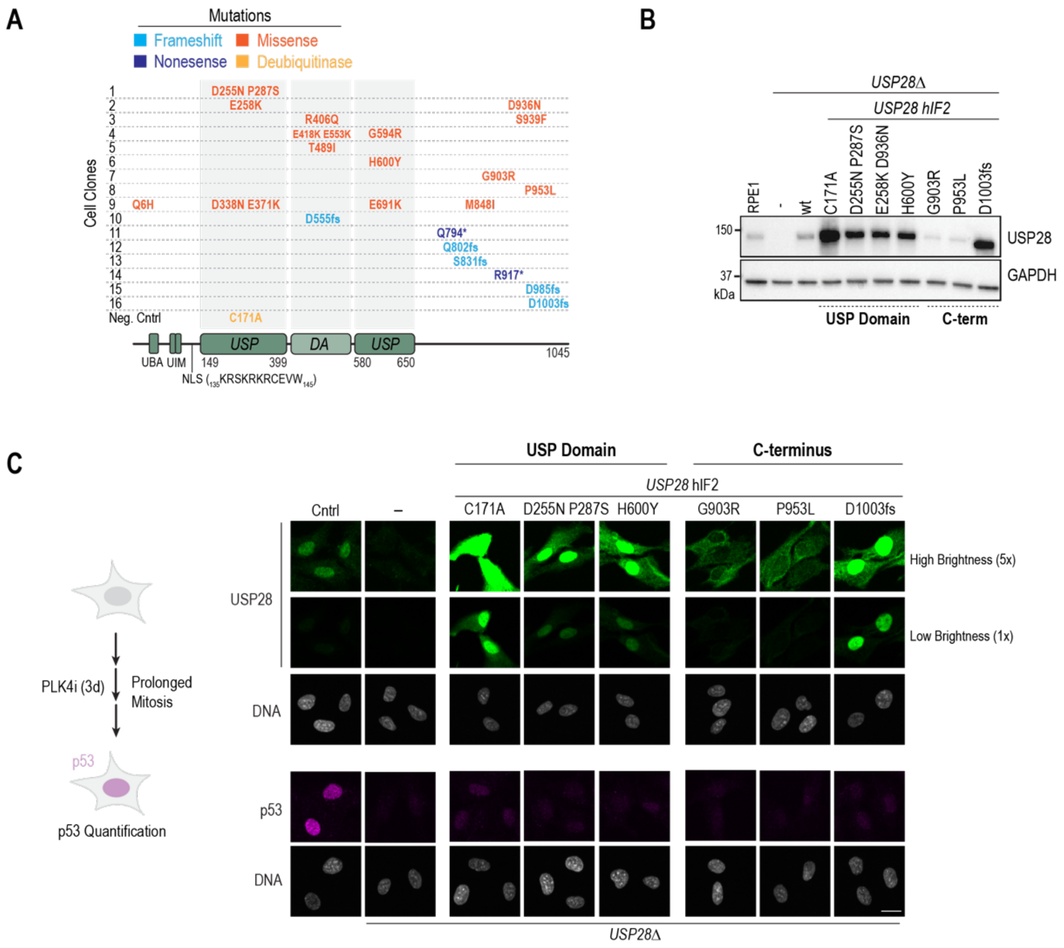


**D**

**USP28hIF2**

**Wildtype** *deleted bases*

CTA CCC AGA CTT CTA GAT CCT TCT GCA GAA ATC ATC GTC TTG AAA GAG ... ACA GTG AAA TAA
L P R L L D P S A E I I V L K E T V K *
998 1003 1013 1045

**Clone 16: D1003fs** *Premature stop codon*

CTA CCC AGA CTT CTA GAG AAA TCA TCG TCT TGA
L P R L L E K S S S *
998 1007

**Figure S3: Characterization of spontaneous mutations in USP28**

**(A)** Cell clones expressing *USP28*$^{hIF2}$ with spontaneous mutations. The locations of clone-specific mutations within the USP28 gene are illustrated. The generation of these clones is depicted in Figure 2A. A summary is shown in Figure 2B.

**(B)** Cell lysates of mitotically arrested cells (Nocodazole, 100 ng/ml, 16 h) with the indicated genotype. The expression of wildtype and mutant USP28 is shown for comparison. GAPDH served as a loading control.

**(C)** Visualization of the USP28 expression and localization in control, *USP28*Δ and *USP28*$^{hIF2}$ mutant transgene expressing *USP28*Δ RPE1 cells (see Figure 2D for comparison). Visualization of p53 expression and stability following three days of PLK4 inhibition to prolong mitosis as indicated in the left schematic. The nucleus was visualized with the DNA-staining compound Hoechst 33342. Quantification of nuclear USP28 levels is shown in Figure 2D. Scale bar: 10 µm.

**(D)** Illustration of the genetic alteration of clone 16 (D1003fs), which has a 10 base deletion at codon 1003 leading to a premature stop codon at position 1008.

Figure S4

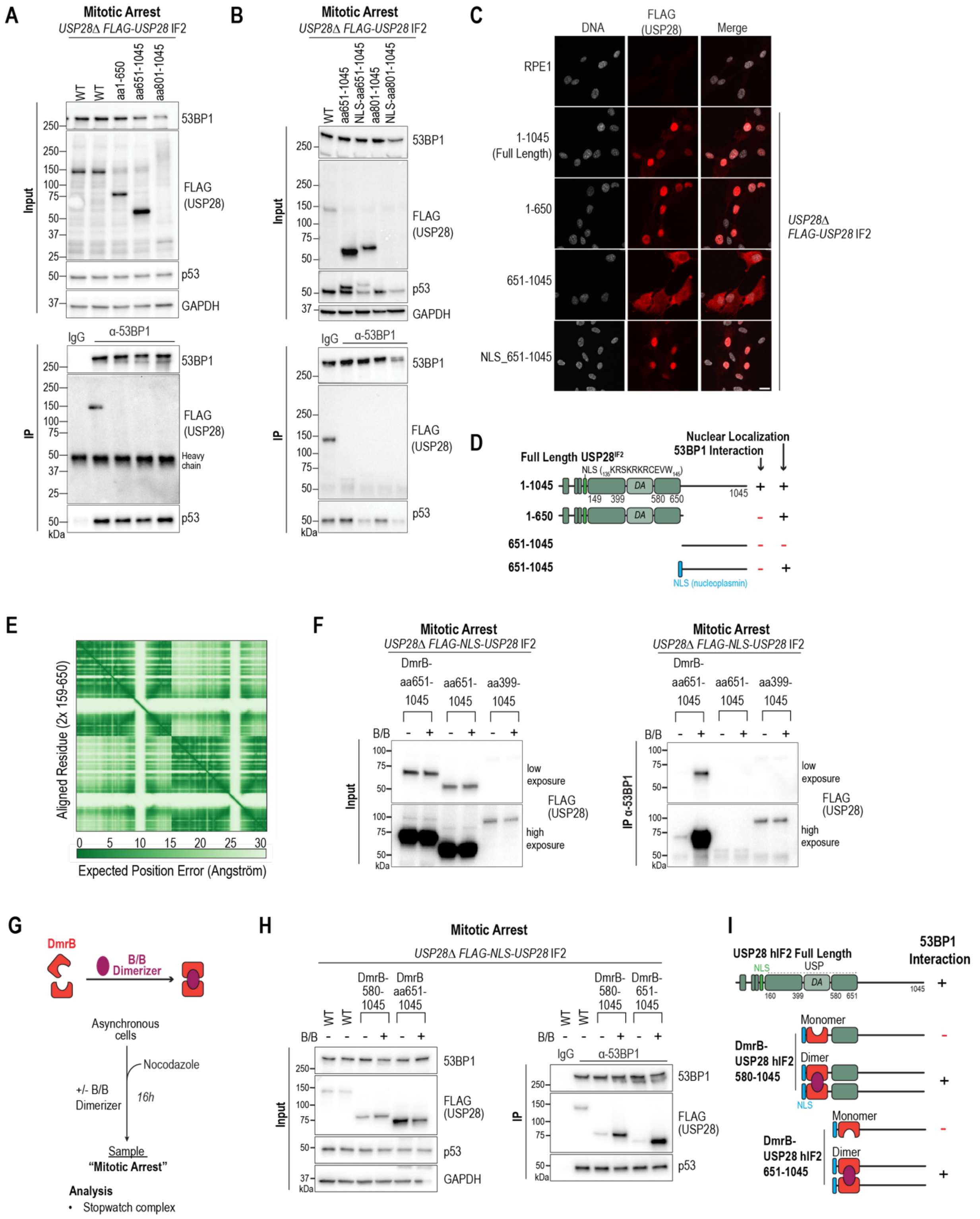

A
Mitotic Arrest
USP28Δ FLAG-USP28 IF2
WT
WT
aa1-650
aa651-1045
aa801-1045
Input
53BP1
FLAG (USP28)
p53
GAPDH
IgG
α-53BP1
IP
Heavy chain
kDa
B
Mitotic Arrest
USP28Δ FLAG-USP28 IF2
WT
aa651-1045
NLS-aa651-1045
aa801-1045
NLS-aa801-1045
C
DNA
FLAG (USP28)
Merge
RPE1
1-1045 (Full Length)
1-650
651-1045
NLS_651-1045
USP28Δ FLAG-USP28 IF2
D
Nuclear Localization
53BP1 Interaction
Full Length USP28IF2
NLS (136KRSKRKRCEVW146)
DA
1-1045
1-650
651-1045
651-1045
NLS (nucleoplasmin)
E
Aligned Residue (2x 159-650)
Expected Position Error (Angström)
F
Mitotic Arrest
USP28Δ FLAG-NLS-USP28 IF2
DmrB-aa651-1045
aa651-1045
aa399-1045
B/B
low exposure
high exposure
FLAG (USP28)
IP α-53BP1
G
DmrB
B/B Dimerizer
Asynchronous cells
Nocodazole
+/- B/B Dimerizer
16h
Sample
"Mitotic Arrest"
Analysis
• Stopwatch complex
H
Mitotic Arrest
USP28Δ FLAG-NLS-USP28 IF2
DmrB-580-1045
DmrB aa651-1045
DmrB-651-1045
I
USP28 hIF2 Full Length
53BP1 Interaction
NLS
USP
Monomer
Dimer
DmrB-USP28 hIF2 580-1045
DmrB-USP28 hIF2 651-1045

**Figure S4: Characterization of USP28 domains that are crucial for the interaction with 53BP1**

**(A-B)** Analysis of 53BP1 immunoprecipitates to determine complex formation with USP28 fragments in mitotically arrested cells (Nocodazole, 100 ng/ml, 16 h). The USP28 fragment 801-1045 was not stably expressed. (B) The USP28 fragment 651-1045 was fused to an NLS from nucleoplasmin. Inputs are soluble supernatants. IP, immunoprecipitate. GAPDH served as a loading control.

**(C)** Subcellular localization of USP28 fragments with and without the NLS, showing the influence of NLS on nuclear targeting. Scale bar: 10 μm.

**(D)** Overview of tested USP28 fragments in (A-C) and their capacity to localize in the nucleus and interact with 53BP1 in mitotically arrested cells. Endogenous and nucleoplasmin NLS are shown in green and blue, respectively.

**(E)** AlphaFold-predicted aligned error map for the USP28 dimer structure, indicating regions of varying confidence in the predicted structural model.

**(F)** Immunoblot showing expression levels of USP28 fragments under different exposure conditions to illustrate relative abundance. The USP28 blotting data from the same membrane with two different exposure times is shown in Figure 4I.

**(G)** Schematic of an inducible dimerization domain. Asynchronous cells were treated for 16h with Nocodazole (100 ng/ml) and the B/B dimerizer (100 nM) to induce dimerization. Mitotic cells were harvested and analyzed by immunoprecipitation.

**(H)** Immunoprecipitation analysis showing that USP28 C-terminal fragments (amino acids 580-1045 and 651-1045) form a complex with 53BP1 as a dimer induced by DmrB in the presence of B/B Dimerizer but not as a monomer in the absence of the B/B Dimerizer, indicating dimerization-dependent interaction. See also Figure 4H-J.

**(I)** Summary of (H) illustrating the requirement of USP28 dimerization for effective C-terminal interaction with 53BP1.

Figure S5

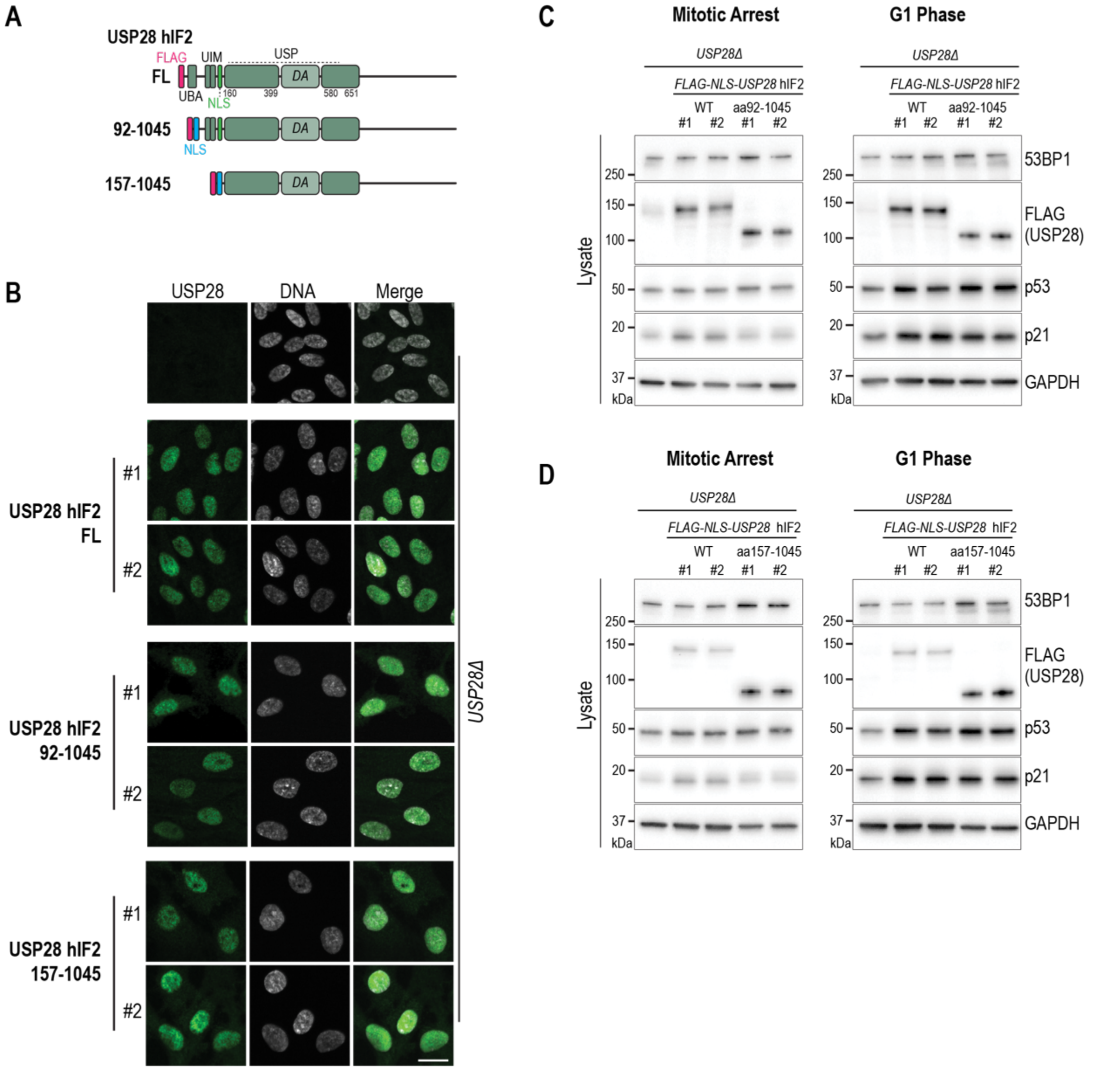


**Figure S5: UBA and UIM are dispensable for the response to prolonged mitosis**

**(A)** Schematic representation of the full-length USP28 (FL) and two N-terminally truncated fragments that lack the UBA (92-1045) or the UBA and UIM (157-1045). All transgenes are fused to a FLAG tag for comparison.

**(B)** Immunostaining of USP28 transgenes (depicted in A) are expressed in *USP28*Δ RPE1 cells. Transgenes are fused to a FLAG tag and NLS from nucleoplasmin, enabling visualization and nuclear localization. Representative images of two independent single clones are shown for each transgene. Scale bar: 10 µm

**(C-D)** Immunoblot analyses of USP28 transgenes (depicted in A), assessing their ability to stabilize and activate p53 following release from prolonged mitotic arrest. Lysates were generated as described in Figure 1F. Lysates are soluble supernatants. GAPDH served as a loading control.

Figure S6

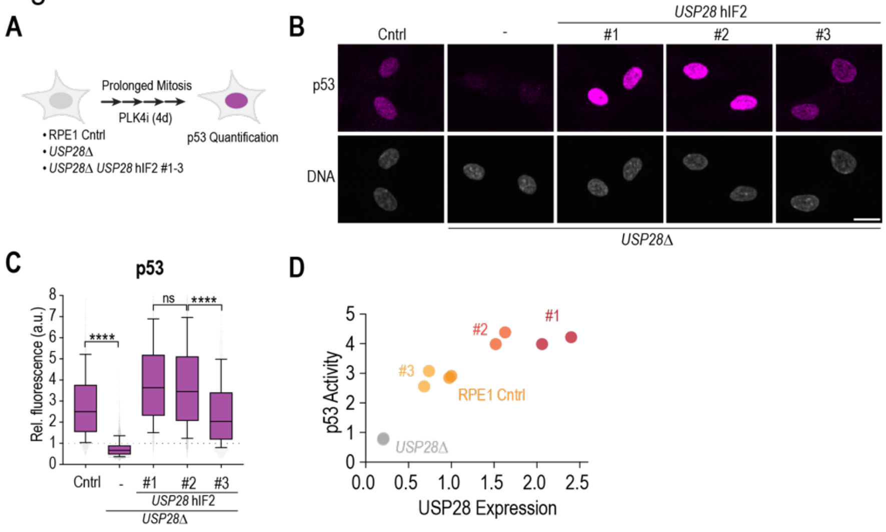


**Figure S6: USP28-dependent sensitivity to prolonged mitosis**

**(A)** Illustration of the method used to assess sensitivity to prolonged mitosis. Cells are fixed and stained for p53 stabilization following a four-day incubation with the PLK4 inhibitor Centrinone, which induces prolonged mitoses. See also Figure 6E-G to compare with a three-day incubation with the PLK4 inhibitor.

**(B)** Immunostaining of RPE1 control cells, *USP28*Δ cells, and the *USP28*$^{\mathrm{hIF2}}$ expressing clones for p53 following prolonged mitosis induced by PLK4 inhibition. The nucleus was visualized with the DNA-staining compound Hoechst 33342. Scale bar: 10 µm.

**(C)** Quantification of immunostaining results from (B). Data is represented as box-and-whisker plots. Mean and 10-90 percentile (error bars) are indicated. Data is normalized to USP28 expression in RPE1 cells (dotted line at 1). Statistical significance was assessed using one-way ANOVA (**$P < 0.01$; ***$P < 0.001$; ****$P < 0.0001$). Sample size (n) > 2,500 cells per condition.

**(D)** Graph showing the relationship between USP28 expression levels and p53 stabilization after 4 days of PLK4 inhibition-induced prolonged mitosis. Each data point represents a replicate for the indicated cell lines.
